# Hierarchical decoding of targeting tripeptide motif by the cytosolic iron-sulfur cluster assembly targeting complex

**DOI:** 10.64898/2026.04.06.716496

**Authors:** Anastasiya Buzuk, Omeir Khan, Soyoon Kang, Leah Yim, Sandor Vajda, Deborah L Perlstein

## Abstract

Iron-sulfur (Fe-S) clusters are essential cofactors required for diverse cellular processes, yet how the Fe–S cluster biogenesis machinery selectively recognizes apo-client proteins remain poorly understood. In eukaryotes, many cytosolic and nuclear Fe–S proteins are recruited to the cytosolic iron-sulfur cluster assembly (CIA) system through a short C-terminal targeting complex recognition (TCR) motif having a [ILM]-[DES]-FW] consensus. Currently, the physicochemical properties underlying this molecular recognition event are undefined. By combining quantitative binding measurements, bioinformatic analysis, and structural modeling, we define the molecular basis for TCR peptide recognition by the CIA targeting complex (CTC). This systematic energetic dissection reveals a hierarchy of binding determinants, in which the side chain and C-terminal carboxylate of the aromatic residue provide the dominant energetic contributor, whereas the upstream residues modulate affinity in a sequence context-dependent manner. Computational docking and molecular dynamics simulations identify an interfacial binding site at the Cia1-Cia2 interface that can accommodate these TCR moieties complementary interaction surfaces. Mutational analysis the identified interaction site is consistent with an aromatic pocket and an adjacent hydrophobic groove on Cia2 accommodating the TCR’s terminal aromatic and antepenultimate aliphatic residues. Together, these results reveal the physicochemical decoding grammar by which the CTC recognizes targeting peptides with divergent sequences, illustrating how short targeting motifs can achieve both the specificity and adaptability required for Fe–S protein maturation.

**Significance Statement:** Iron-sulfur (Fe-S) clusters are essential metallocofactors requiring dedicated pathways for their assembly and insertion into proteins. How Fe–S cluster biogenesis systems selectively recognize their Fe-S binding client proteins in the crowded cellular environment remains poorly understood, in part because these machineries must engage dozens of clients rather than relying on the one-to-one metallochaperone-client pairings commonly used to assemble other types of metalloproteins. In eukaryotes, many cytosolic and nuclear Fe–S proteins are recruited to the cytosolic iron–sulfur assembly (CIA) machinery through a C-terminal targeting tripeptide motif. Here we combine biochemical measurements and structural modeling to define the molecular rules governing recognition of CIA targeting peptides. We show that a hierarchy of physicochemical peptide features, rather than a strict sequence consensus, guides recruitment of clients to the targeting complex, explaining how a single binding site can decode multiple targeting peptide signals.

## Main text

Iron-sulfur (Fe-S) cofactors are among the most ancient and versatile cofactors in biology, enabling essential processes ranging from energy conversion to genome maintenance. Because Fe–S clusters do not spontaneously assemble in vivo, all organisms rely on dedicated biosynthetic pathways to build and deliver these metallocofactors to apo-client proteins.(1-4) A central challenge for Fe–S maturation systems or any pathway acting on multiple substrates is specificity: how does a targeting apparatus recognize the correct client in a complex and crowded cellular milieu?(5-7) Most metalloprotein maturation systems address this specificity problem through dedicated metallochaperone-client pairs. In contrast, Fe–S biogenesis pathways must recognize dozens of structurally and functionally diverse clients through a shared set of targeting factors, raising the question of how such systems achieve selective yet adaptable client recognition.

Recognition of eukaryotic cytosolic and nuclear Fe–S proteins is mediated by the cytosolic iron-sulfur cluster assembly (CIA) targeting complex (CTC; **Fig. 1A**).(8-11) Approximately one quarter of the >30 known CIA clients are recruited to the CTC through a C-terminal targeting complex recognition (TCR) motif.(11, 12) This tripeptide motif functions analogously to other short linear motifs (SLiMs) that direct protein localization, modification, and complex formation, providing a compact and universal signal for directing various biosynthetic machineries to their clients.(13) Identification of the [ILM]-[DES]-[FW]-COOH targeting motif provided an initial framework for deciphering the CIA targeting mechanism, yet the molecular principles by which the CTC interprets this motif remain poorly defined. A putative binding site at the interface of Cia1 and Cia2 has been proposed, yet there is no structural information about how the TCR engages the targeting complex and limited biochemical evidence defining the energetic contributions of individual residues within the motif (**Fig 1B**).(12) Therefore, it remains unclear how this short peptide signal achieves both the specificity and tolerance for the sequence variability amongst CIA clients.

**Figure 1.**
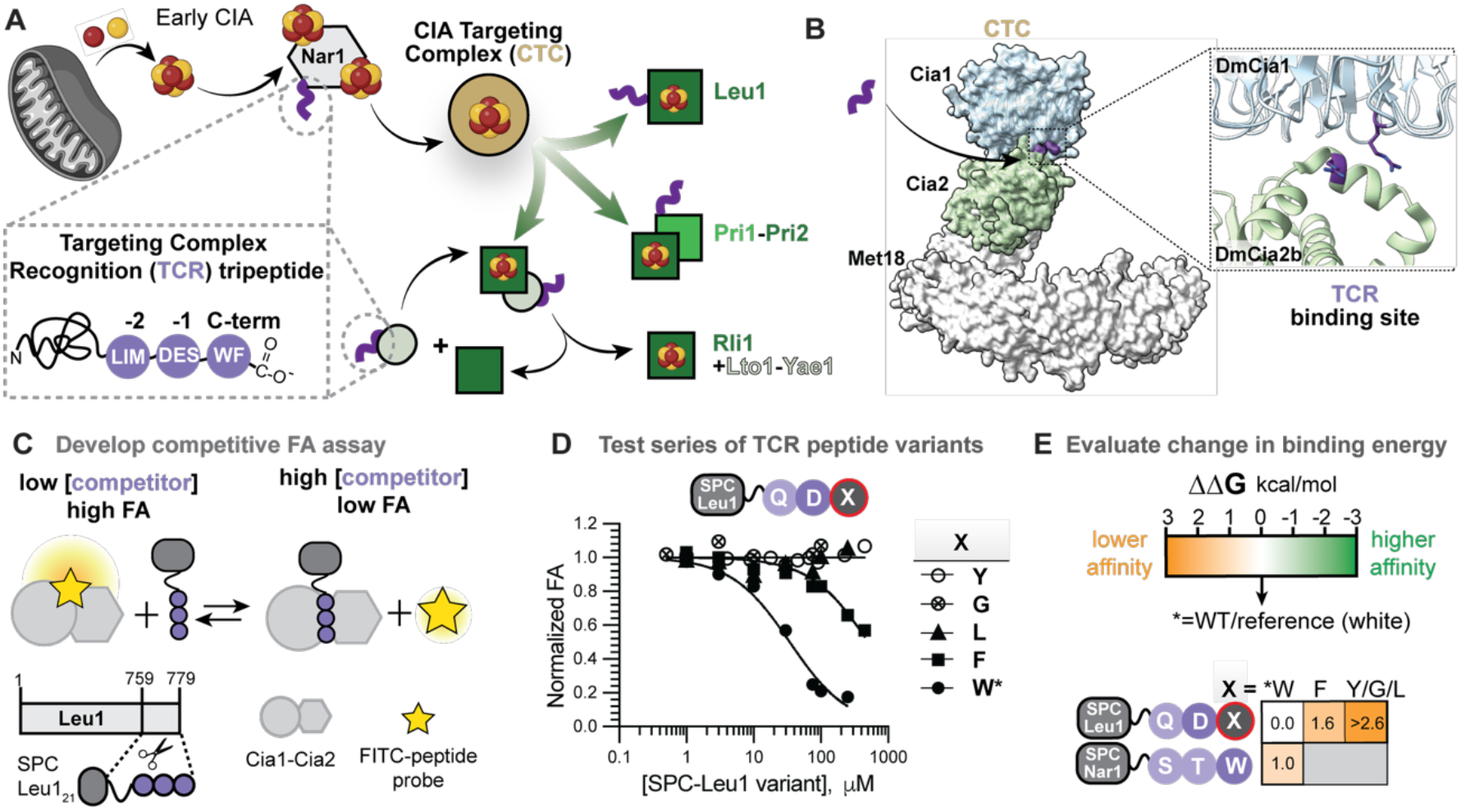
Strategy to dissect molecular recognition of the TCR tripeptide by the Cia1-Cia2 complex. **(A)** In the late stages of the CIA pathway, the CIA targeting complex (CTC, gold) receives an Fe-S cluster from Nar1 and delivers it to client proteins (green), a subset of which are recruited via their targeting complex recognition (TCR, purple) tripeptide. Different modes of CTC recruitment via a TCR motif are illustrated. **(B)** The CTC comprises Cia1, Cia2, and Met18 (yeast nomenclature; PDB: 6TC0). Two conserved Arg residues (purple sticks, *D. melanogaster* numbering) at the Cia1-Cia2 interface have been implicated in TCR peptide binding. **(C)** Schematic of the fluorescence anisotropy (FA) assay used to quantify binding of TCR peptide variants. **(D)** Representative FA competition curves for the indicated SPC-Leu1 variants. **(E)** Changes in binding free energy (ΔΔG) relative to the WT reference (*) are color-coded orange to green, as indicated. TCR consensus residues are circled in dark purple, and nonconsensus residues are in light purple.

Furthermore, several observations about the CTC’s interaction with the TCR motif remain unexplained and, in some cases, appear contradictory. For example, substitution of the C-terminal tryptophan residue in the benchmark CIA client Leu1 with phenylalanine, swapping one consensus residue for the other, abolishes detectable CTC binding *in vitro* but does not impair Fe–S cluster acquisition *in vivo*.(11) Conversely, replacement of the strongly conserved penultimate acidic residue with a nonconsensus alanine does impact CTC binding to Leu1 or cofactor maturation efficiency.(11) The functional role of the antepenultimate aliphatic residue is even less clear, as prior studies focused on substituting the Leu1’s nonconsensus glutamine residue with another nonconsensus residue, limiting interpretation of the reported lack of effect.(11).

Such inconsistencies are characteristic of SLiM-mediated interactions, in which degenerate sequence patterns often mask the underlying physicochemical determinants of binding.(13, 14) In many SLiM recognition systems, multiple peptide ligands engage a common binding pocket through partially shared interaction features rather than strict sequence conservation.(15) These common SLiM properties have frustrated efforts to predict targeting motifs from sequence alone, as it remains unclear how roughly 75% of CIA clients are recruited to the CTC.(14) Moreover, modern structure-function prediction methods can provide unprecedented insights into the architecture of macromolecular assemblies, but they often struggle to accurately model peptide-protein interfaces involving such short, flexible, and degenerate motifs.(16, 17) These challenges highlight the need for quantitative energetic dissection and structural analysis of TCR peptide-based recruitment to the targeting complex and to define the molecular grammar driving SLiM peptide decoding in the CIA system.

To address these challenges, we combined quantitative binding measurements, bioinformatic analyses, and structural modeling to dissect the molecular determinants governing recognition of TCR peptides by the CIA targeting complex. Our results reveal a hierarchy of binding determinants in which both the side chain and C-terminal carboxylate of the terminal aromatic residue provide the dominant energetic contributions. We further demonstrate that the upstream residues modulate binding affinity in a sequence-context-dependent manner. Computational modeling identifies a peptide binding site at the interface of the Cia1 and Cia2 subunits of the targeting complex consistent with constraints derived from biochemical experiments, revealing the complementary interaction surfaces that engage the targeting peptide. Together, these findings define a physicochemical decoding grammar for TCR peptide recognition, resolving prior discrepancies and providing a framework for understanding how the CIA machinery achieves the selective recruitment of its diverse client proteins.

## Results

### Quantitative platform for dissecting TCR peptide decoding by the CIA targeting complex

To define how the CIA targeting complex decodes the targeting complex recognition (TCR) peptide, we developed a quantitative fluorescence anisotropy (FA)-based competitive binding assay. In this format, unlabeled TCR peptide constructs compete with FITC-labeled TCR peptide probes, such as His-Gln-Asp-Trp (HQDW; probe derived of *Saccharomyces cerevisiae* (*Sc*) Leu1) for binding to the *Chaetomium thermophilum* (*Ct*) Cia1-Cia2 complex (**Fig. 1C**), selected for its thermal stability which increases assay reproducibility. The targeting complex subunit Met18 was omitted because it is dispensable for TCR peptide recognition.(12) To dissect the intrinsic contribution of individual targeting peptide residues without synthesizing a large number of FITC-labeled peptides, we displayed TCR peptide variants as C-terminal fusions to a SUMO peptide carrier (SPC), a previously validated scaffold for probing CTC-TCR interaction.(11, 12) Whereas prior studies used SPC constructs in qualitative interactions assays, here we integrate this display technology into a quantitative competitive assay format, allowing for evaluation of a large panel of peptide variants under identical conditions. The resulting competitive binding assay provides a robust quantitative platform for comparing the intrinsic energetic contributions of individual targeting peptide moieties, enabling a residue-level dissection of the recognition energetics underlying TCR peptide decoding (**Fig. 1D** and **E**).

We first asked whether residues upstream of the TCR tripeptide measurably contribute to the CTC’s recognition of client C-termini. Short peptides corresponding to the final three or final 21 residues of the *Sc*Leu1 C-terminus were fused to the SPC scaffold to generate two SPC-Leu1 constructs. Both competitively displaced a FITC-tetrapeptide probe from *Ct*Cia1-Cia2 complex with indistinguishable affinities (**Fig. S1A, Table S1**), establishing that the final three amino acids of Leu1 capture all the measurable binding energy derived from its C-terminal tail.

Replacement of the Leu1 21-mer of SPC-Leu1 with a 10-mer derived *Sc*Nar1 resulted in ∼5-fold weaker binding (**Fig. S1, Table S1**), consistent with the *Sc*Nar1 TCR motif (Ser-Thr-Trp) matching the consensus at only one position versus two for the *Sc*Leu1 motif (Gln-Asp-Trp). Importantly, the peptide presentation mode did not measurably impact *Sc*Nar1’s TCR binding affinity, as replacement of the SPC with SNAP or removal of the carrier altogether yielded comparable *Sc*Nar1 TCR peptide binding affinities (**Fig. S1B**). In sum, development of the competitive peptide binding assay revealed that TCR peptide recognition is encoded within the final few residues of client proteins, and that the measured binding affinities of the SPC-presented peptides reflect the intrinsic properties of the targeting peptide rather than its mode of presentation.

### The C-terminal aromatic residue is a primary energetic binding determinant for TCR peptide recognition

Having validated the competitive binding assay, we applied it to reveal the energetic contribution of the C-terminal aromatic residue to CTC binding, using variants of SPC-Leu1. Substitution of the C-terminal tryptophan with phenylalanine resulted in a 16-fold increase in the IC_50_, indicating substantially reduced affinity results from even this seemingly conservative change involving swapping one consensus residue for another (**Fig. 1D** and **E, Table S1**). Substitution of Trp with Tyr, Leu, or Gly abolished detectable binding, corresponding to losses of at least two orders of magnitude in affinity relative to variants ending in tryptophan or phenylalanine. Importantly, these data reveal a hierarchy in binding affinity that could not have been inferred from consensus sequence alone, as tryptophan and phenylalanine are equally represented at the C-termini of CIA clients but bind with an order of magnitude different affinities.(11, 12)

To determine whether the reduced affinity observed upon replacing the C-terminal Trp with Phe reflects intrinsic differences in binding energy rather than secondary effects such as peptide conformation, we tested the binding of individual N-acetylated amino acids. N-acetyl-Trp bound *Ct*Cia1-Cia2 with a *K*_*D*_ value ≥630 µM, only 13-fold weaker than that of SPC-Leu1 (**Fig. 2A, Fig. S2A, Table S2**). The binding of N-acetyl-Phe was weaker (*K*_*D*_ ≥10 mM), and N-acetyl-Tyr showed no detectable binding, mirroring the hierarchy observed when these residues were positioned at the C-terminus of the TCR tripeptides. These measurements provide direct experimental evidence that the terminal aromatic comprises a dominant TCR peptide binding determinant, and that the observed hierarchy arises from differences in intrinsic binding energy arising from side chain interactions rather than its sequence-context. This hierarchy additionally explains why binding of a Phe-terminated targeting peptide could not be detected *in vitro* yet it mediated client Fe–S maturation *in vivo*,(11) its reduced binding affinity likely falls below the detection threshold of the qualitative pull-down assay.

**Figure 2.**
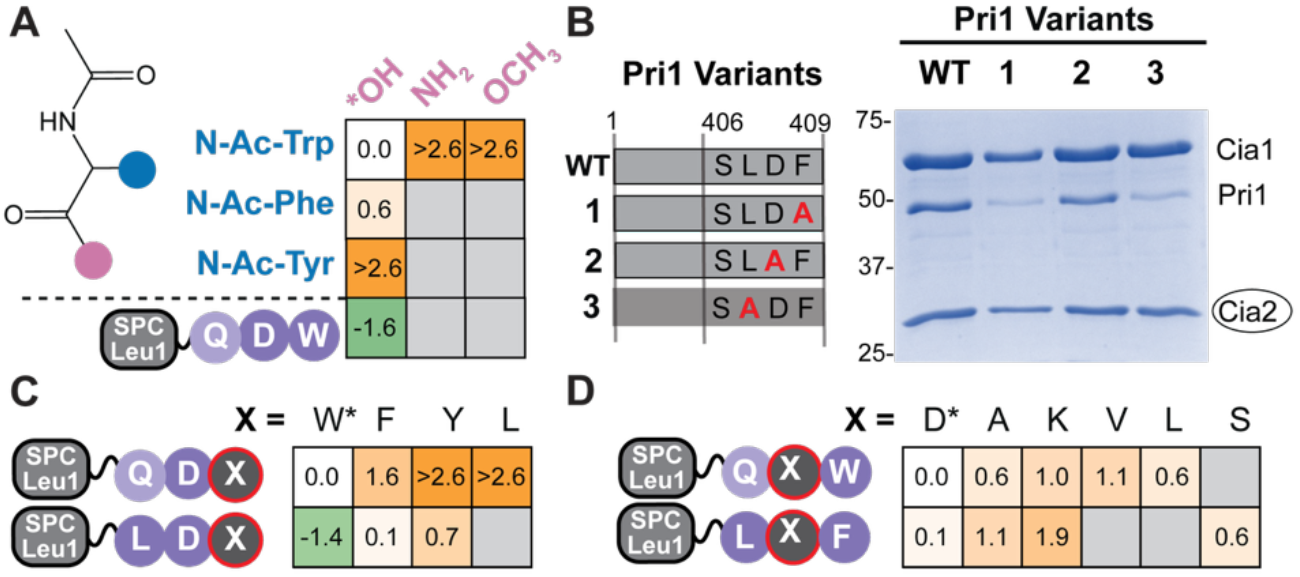
Energetic dissection of TCR peptide binding determinants. (**A**) Changes in binding free energy (ΔΔG) of SPC-Leu1 and N-acetylated (N-Ac) amino acids relative to N-acetyl-tryptophan reveal the contributions from the aromatic side chain (blue) and main-chain carboxylate (pink). Fluorescence anisotropy (FA) data are in **Fig S2**. (**B**) Affinity copurification of wild-type (WT) *Sc*Pri1 or the indicated TCR motif variants with *Ct*Cia1-Cia2 (strep-tagged bait, circled), analyzed by SDS-PAGE followed by Coomassie staining. Substitution of the antepenultimate or terminal residue of Pri1 disrupts its interaction with the targeting complex. Uncropped gels including input samples and controls are in **Fig. S3**. (**C-D**) ΔΔG heatmaps illustrating how substitution at the -2 position (Gln→Leu; C) or -1 position (D) alters the intrinsic binding energy relative to that of SPC-Leu1 with Gln-Asp-Trp terminal tripeptide. Primary data are in **Fig. S4D**.

### An aliphatic antepenultimate residue compensates for weakened binding of phenylalanine-terminated TCR peptides

Given the reduced affinity of Phe-terminated motifs, we asked how such naturally occurring targeting peptides compensate for this energetic penalty. To probe this question, we examined the TCR motif of the yeast primase small subunit of primase (*Sc*Pri1), which ends in a Leu-Asp-Phe targeting peptide proposed to facilitate Fe–S cluster maturation of the primase Pri2 subunit (**Fig. 1B**).(10, 11) *Sc*Pri1 copurified with *Ct*Cia1-Cia2 in a manner dependent on its terminal Phe residue (**Fig. 2B, Fig. S3A**). Additionally, Arg163 of *Ct*Cia2, a key TCR peptide binding hotspot residue,(12) was required for this interaction (**Fig. S3B**), confirming that this Phe-terminated sequence binds at the same site as targeting motifs ending in tryptophan. Strikingly, substitution of *Sc*Pri1’s antepenultimate leucine residue (L407A) abrogated interaction with *Ct*Cia1-Cia2 (**Fig. 2B, Fig. S3A**), suggesting that the -2 residue may compensate for the reduced binding affinity of Phe-terminated TCR motifs.

Motivated by the importance of Pri1’s leucine residue for CTC interaction, we introduced a glutamine-to-leucine substitution into FITC-labeled tetrapeptide probes and measured the resulting impact on binding to *Ct*Cia1-Cia2. This substitution increased binding energy by 1.3 kcal/mol relative to FITC-HQDW (**Fig. S4A, Table S3**). Introduction of both a -2 Leu residue and a terminal Phe residue, generating a Leu-Asp-Phe peptide, resulted in comparable binding affinity as the Gln-Asp-Trp peptide regardless of its presentation as a FITC-labeled peptide or using the peptide carrier (**Fig. 2C, Fig. S4, Table S3, Table S4**). Control experiments confirmed that TCR peptides with antepenultimate leucine or glutamine residues bind at the same site on *Ct*Cia1-Cia2 (**Fig. S4**). We concluded that a leucine residue at the -2 position contributes directly to the intrinsic binding energy of TCR peptides, providing the mechanistic basis for enrichment of an aliphatic residue at the antepenultimate position of the TCR motif revealed though bioinformatic analysis.

Having established that a -2 leucine residue can compensate for the reduced binding affinity resulting from a tryptophan-to-phenylalanine substitution, we next asked if it could also facilitate binding of TCR peptides ending in tyrosine or aliphatic residues. Although we could not detect binding of N-acetyl Try or SPC-Leu1 ending in Gln-Asp-Tyr, an SPC-presented Leu-Asp-Tyr tripeptide bound to *Ct*Cia1-Cia2 with only a modest ∼3-fold reduction in binding affinity relative to wild-type SPC-Leu1 (**Fig. 2C, Fig. S4D, Table S4**), but we could not detect binding of Leu-terminated constructs. We concluded that a -2 leucine residue can compensate for the reduced affinity of tyrosine-terminated TCR peptides, consistent with our observations with Phe-terminated peptides, and functions more broadly to tune binding energy when lower-affinity residues are presented at the other positions.

### The C-terminal carboxylate is essential for TCR recognition, whereas the penultimate acidic residue is dispensable

Having defined the roles of the hydrophobic moieties of the TCR motif, we turned to defining the contributions of the TCR peptide C-terminus and the acidic (-1) side chain (**Fig. 1A**, dashed box). Although the TCR peptide-binding site is enriched in basic residues, suggesting that electrostatic interactions with negatively charged groups could contribute to recognition, this interpretation is difficult to reconcile with the observation that the penultimate Asp residue of full-length Leu1 is dispensable for CTC recruitment *in vitro* and for efficient maturation of Leu1’s Fe–S cluster *in vivo*.(11, 12) To resolve this apparent discrepancy further define the TCR peptide decoding grammar, we systematically varied each acidic moiety of the TCR motif to define its contribution to CTC recruitment.

To probe the contribution of the terminal carboxylate, we compared the binding of N-acetyl Trp with derivatives in which the carboxylate was replaced with a carboxamide or a methyl ester. Neither derivative displaced the FITC-HLDW probe from *Ct*Cia1-Cia2 (**Fig. 2A, Fig. S5A**), indicating that the negative charge is critical for interaction. To test the role of the C-terminus in the context of a full TCR peptide, we appended an Asp-Ser-Gly tripeptide to SPC-Leu1, effectively relocating the negative charge of the carboxy terminus to an aspartate side chain. This modification also strongly disrupted binding of the highest affinity TCR tripeptide (Leu-Asp-Trp, **Fig. S5B**). These results establish the main-chain carboxylate as an essential and non-substitutable TCR peptide-binding determinant, indicating both its negative charge and precise positioning are required for interaction. Even minimal perturbations that neutralized or relocated the negative charge abolished detectable binding.

In contrast, substitution of the penultimate acidic residue had surprisingly small effects on binding. Its replacement with aliphatic residues (Ala, Val, Leu) resulted in relatively modest changes in binding affinity (**Fig. 2D, Fig. S6, Table S5**), explaining why the Asp-to-Ala substitution in full-length Leu1 had little impact on TCR mediated interactions *in vitro* or *in vivo* despite strong conservation of the penultimate aspartate residue across fungal orthologs.(11) Together these data indicate that conservation of the penultimate acidic residue cannot be explained through a simple model and suggested that its contribution to CTC recruitment could be dependent on the sequence context.

To reveal potential sequence-context dependent constraints, we revisited the TCR consensus analysis, analyzing 699 C-terminal tripeptides from plants, animals and fungi (**Fig. S7**). Based on the 10-fold difference in binding affinity we observed for Trp- and Phe-terminated TCR peptides, we hypothesized the C-terminal residue could influence sequence preference at the -1 and -2 positions. Therefore, we analyzed Phe-terminated and Trp-terminated sequences separately. Among the 88 Trp-terminated TCR peptides, 37.5% contained an Asp (4.3-fold enriched) or Glu (2.4-fold enriched) at the -1 position. Unexpectedly, Lys was the next most frequent residue, present in 13.6% of sequences, corresponding to a 2.4-fold enrichment relative to its frequency in the human proteome.

In contrast, the 87 Phe-terminated motifs exhibited a stronger preference for a penultimate acidic residue, with 55.2% containing Asp (8.5-fold enriched) or Glu (2.1-fold enriched). Consistent with previous bioinformatic analyses,(11) serine was the only other amino acid found in ≥ 10% of the Phe-terminated sequences. Notably, none of the Phe-terminated sequences contained lysine at the -1 position (**Fig. S7**), distinguishing them from the Trp-terminated motifs. This strict exclusion observed exclusively in Phe-terminated TCR sequences suggests an increased reliance penultimate residue when the C-terminal aromatic reside contributes less intrinsic binding energy.

To directly test this prediction, we compared the effects of an Asp-to-Lys substitution in different contexts. When introduced in the Leu1 TCR tripeptide (Gln-Asp-Trp), this substitution caused only a modest ∼5-fold decrease in binding affinity, whereas introduction of lysine into the Pri1 TCR motif (Leu-Asp-Phe) resulted in a much larger ∼20-fold reduction (**Fig. 2D, Fig. S6, Table S5**). These data reconcile the seemingly contradictory observations to reveal a previously overlooked sequence-context dependence. The penultimate acidic residue of the TCR motif is not a universal binding determinant. Instead, it fine-tunes affinity in a sequence-context dependent manner, contributing more strongly to the peptide’s intrinsic binding energy in Phe-terminated motifs as compared to those ending in Trp. In contrast, the terminal carboxylate functions as a universal and non-substitutable binding determinant.

### Computational modeling reveals candidate TCR peptide binding mode

To gain structural insight into TCR peptide recognition, we turned to computational modeling, employing a series of complementary computational approaches guided by the experimentally defined constraints to identify potential TCR peptide binding sites. Focusing on the region of the Cia1-Cia2 interface previously implicated in recruitment of TCR peptides,(12) we first applied E-Map, a structure-guided method that detects ligand-binding hotspots by docking a diverse set of small molecule probes onto a protein surface and identifying regions of probe clustering.(18, 19) The highest-ranking hotspot contained clusters of both apolar and aromatic pharmacophores (**Fig. 3A**), consistent with a site that could accommodate an aromatic side chain. A second prominent hotspot was enriched in hydrogen-bond acceptor pharmacophores, suggestive of a region that could engage the negatively charged moieties of the TCR peptide. Notably, this second cluster was close to an arginine residue previously implicated in TCR peptide binding (**Fig. 3A**).(12) The spatial proximity of these hotspots identifies a candidate region at the Cia1-Cia2 interface that could of simultaneously accommodate the TCR peptide’s dominant binding determinants, the C-terminal aromatic side chain and terminal main-chain carboxylate.

**Fig 3.**
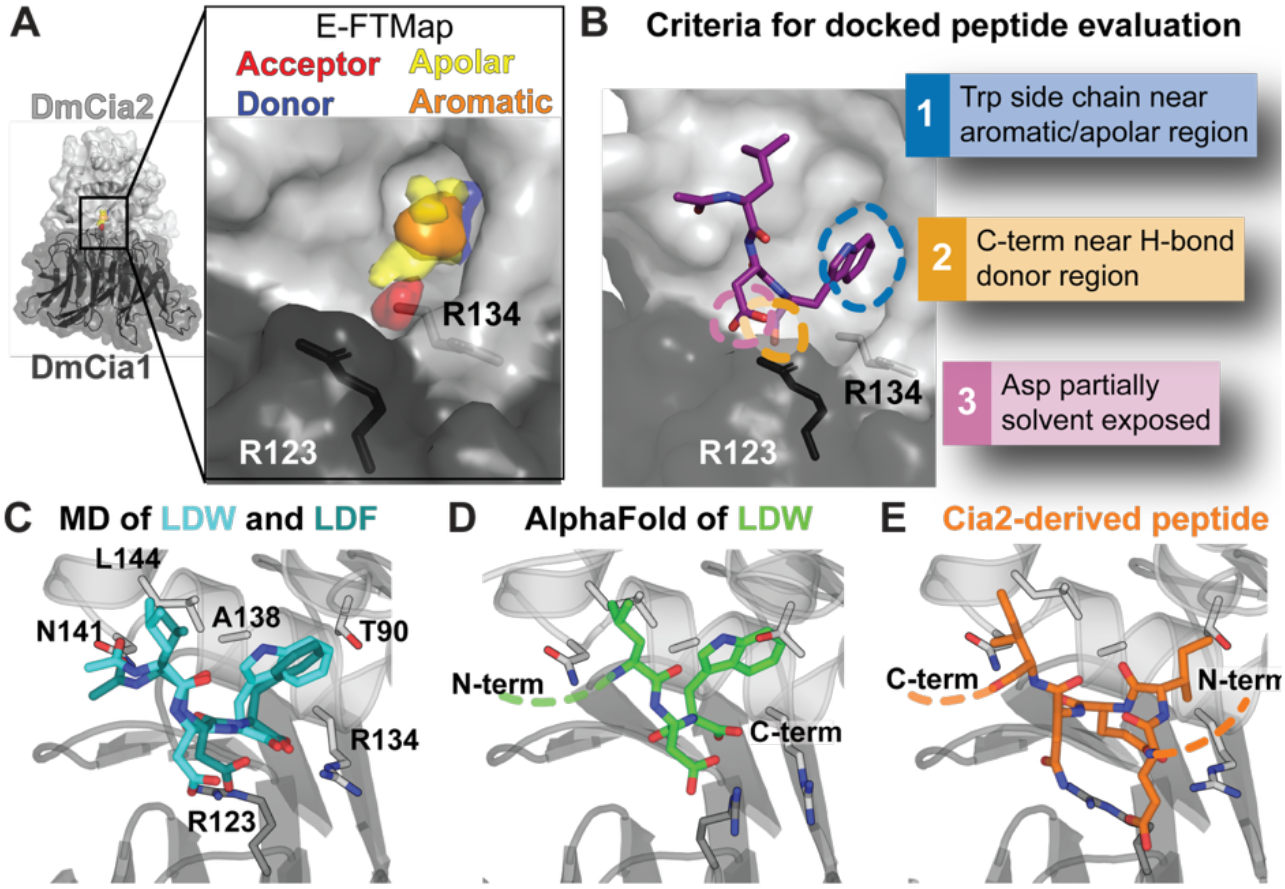
Computational modeling identifies a candidate TCR peptide binding mode. (**A**) E-FTMap analysis of *Dm*Cia1-Cia2b (PDB: 6TBN)(10) identifies hydrogen-bond acceptor (red), hydrogen-bond donor (blue), apolar (yellow), and aromatic (orange) pharmacophore clusters at the Cia1-Cia2 interface, near Arg123 of *Dm*Cia1 and Arg134 of *Dm*Cia2b which have been implicated in TCR peptide binding.(12) (**B**) Docking of the N-acetyl-Leu-Asp-Trp peptide to the Cia1-Cia2 interfacial region. A representative docked pose satisfying experimentally derived constraints is shown. Additional docking solutions are presented in **Fig. S8**. (**C**) Overlay of representative MD-derived binding modes for Leu-Asp-Trp (cyan sticks) and Leu-Asp-Phe (teal sticks) tripeptides illustrating that both can be accommodated within the same interfacial binding region. Residues targeted for substitution to test the computational model are shown as sticks (dark gray, *Dm*Cia1; light gray, *Dm*Cia2b) and positions of N- and C-termini indicated. (**D**) AlphaFold3 prediction of TCR peptide (TTFDKH**LDW**, underlined region shown as green sticks) positions the TCR peptide within the region identified by docking and MD simulations.(17) (**E**) In the crystal structure of the *Dm*Cia1-Cia2 complex,(10) a peptide derived from the N-terminus of *Dm*Cia2b (MPTE**IENI**NPNVY; underlined region shown as orange sticks) is observed in the same interfacial region identified by computational prediction and AlphaFold. The bound peptide displays alternating hydrophobic and acidic residues reminiscent of the physicochemical properties of the TCR motif despite its antiparallel orientation.

Next, we docked a TCR peptide to identify binding mode consistent with both the E-FTMap hotspots and experimentally derived constraints (see **Supporting Text**). An N-acetylated Leu-Asp-Trp tripeptide was therefore docked onto the *Dm*Cia1-Cia2 complex using the FRED docking program and restricting the search to the region surrounding the FTMap-identified hotspots.,(20-23) Docking poses were evaluated using experimentally derived constraints: placement of the C-terminal aromatic side chain within the apolar/aromatic hotspot, formation of favorable electrostatic interactions between the terminal carboxylate and basic residues at the Cia1-Cia2 interface, and partial solvent exposure of the penultimate aspartate residue (see **Supporting Text**). Among the top-ranked docking solutions, one binding mode was identified that satisfied all the criteria (**Fig. 3B, Fig. S8**).

To evaluate the stability of the docked peptide, we performed multiple independent molecular dynamics (MD) simulations of the peptide-Cia1-Cia2 complex (see **Supporting Text**). Across all trajectories, the N-acetyl-Leu-Asp-Trp tripeptide remained associated with the interfacial region and clustering analysis of the resulting peptide conformations revealed a small number of closely related binding modes (**Fig. S9**). Across these binding modes, the tryptophan indole side chain remained positioned within a hydrophobic pocket coincident with the apolar/aromatic hotspot predicted by E-FTMap and the carboxy-terminus and penultimate aspartate side chain remained close to the arginine residues previously shown to be TCR peptide binding hot spot residues.(12) Quantitative analysis of the MD trajectories was consistent with an electrostatic interaction between the peptide C-terminal carboxylate and the guanidinium group of Arg134 of *Dm*Cia2b, with at least one polar contact maintained across >90% of simulation frames (**Fig. S10, Fig. S11**). In contrast, polar interactions involving the penultimate aspartate residue and R123 of *Dm*Cia1 appeared to be more transient, with distances consistent with no polar interaction in 30.5% of the MD frames, aligning with this residue’s modest contribution to the binding free energy of the Leu-Asp-Trp tripeptide. Replacement of the terminal Trp with Phe followed by additional MD simulations yielded comparable binding poses across the most populated clusters (see **Supporting Text, Fig. 3C, Fig. S9**), consistent with the biochemical data showing that the Trp- and Phe-terminated TCR sequences occupy the same binding pocket.

In addition to the anchoring the C-terminal aromatic residue, the MD stimulations suggested a second interaction surface capable of accommodating the antepenultimate aliphatic residue of the TCR motif (**Fig. 3C**). In both Phe and Trp-terminated peptides, the side chain of Leu at -2 position was frequently observed packing against a hydrophobic groove along the surface of Cia2 (**Fig. 3C, Fig. S9**), consistent with a role in augmenting affinity rather than serving as a primary anchor. This groove is lined predominantly by conserved nonpolar residues, suggesting it is optimized to engage aliphatic side chains rather than polar or charged groups. In total, these docking studies combined with MD simulations support a modular recognition mechanism in which distinct features of the TCR motif are independently accommodated by complementary surfaces at the Cia1-Cia2 interfacial region.

To test whether the same TCR peptide binding site could be identified via a different computational approach, we examined AlphaFold3 predictions of the *Dm*Cia1-Cia2 complex bound to a peptide terminating in the highest-affinity TCR sequence (TTFDKHLDW; **Fig. 3D, Fig. S4B**). Across all predicted models, the terminal aromatic and -2 aliphatic residues occupied positions closely aligning with the MD-derived binding poses. Extending this analysis to larger assemblies containing the full-length client proteins terminating in Trp or Phe and the yeast, fly, or human CTC similarly placed the TCR peptide within the same interfacial site, consistent with this binding mode being maintained across species and is compatible with intact CTC-client complexes (**Fig. S13**).

Because accurate prediction of novel peptide-protein interactions remains a challenging regime for structure prediction programs, including AlphaFold,(24, 25) we examined the Cia1-Cia2 structure to identify any overlooked features that could be serving as a structural template for the AlphaFold prediction.(10) To our surprise, we found that an EIENI pentapeptide derived from the N-terminus of Cia2 occupies the same interfacial pocket identified by computational modeling in all structures (**Fig. 3E, Fig. S14A**). Although the pentapeptide adopts an antiparallel orientation relative to that of the TCR motif, its alternating hydrophobic and acidic residues occupy positions corresponding to the aliphatic/aromatic and acidic features of the TCR motif.

To assess functional relevance of the observed interaction of Cia2’s N-terminus with the TCR peptide binding site, we tested both its potential role in regulation client binding via occlusion of the TCR binding site and its ability to compete with the TCR peptide for binding to *Ct*Cia1-Cia2. Truncation of the N-terminus of *Ct*Cia2, thereby removing the pentapeptide, had no detectable effect on binding of FITC-labeled TCR peptide probe (**Fig. S14B**). Likewise, an SPC-presented pentapeptide failed to compete with the TCR probe for binding to *Ct*Cia1-Cia2 (**Fig. S14C**). Nevertheless, the recurrent crystallographic capture of a peptide with TCR-like physicochemical features within the same pocket identified by AlphaFold predictions and computational modeling provides independent structural support for the location of the TCR peptide binding site.

### Biochemical validation of the TCR peptide binding mode

To experimentally test the predicted TCR peptide binding mode, we substituted several conserved residues predicted to contact the peptide (**Fig. 3C**). Across all variants, expression level, purification yield, and *Ct*Cia1-Cia2 complex stability were compared to those of the wild-type complex to identify any substitutions resulting in global disruption of protein folding rather than localized perturbations of peptide binding (**Table S6, Fig. S15**).

The MD-derived peptide binding models predicted that the C-terminal aromatic residue of the TCR motif is accommodated within a hydrophobic pocket formed predominantly by residues derived from the Cia2 subunit (**Fig. 4A**). To test the functional importance of this pocket, we reduced its volume by substituting Ala181 of *Ct*Cia2 (corresponding to Ala138 of *Dm*Cia2b) with the β - branched residue valine (**Fig. 4B**). The resulting *Ct*Cia1-Cia2^A181V^ complex was stable and readily purified but exhibited a pronounced 10-15-fold reduction in TCR peptide binding affinity (**Fig. 4C, Fig. S15A, Table S6**). We also substituted Thr133 of *Ct*Cia2 with alanine, since the corresponding residue in *Dm*Cia2b (Thr90, **Fig. 3C**) appeared to make a polar contact with the indole nitrogen in some of the MD simulation trajectories, but this variant had no measurable impact on FITC-HLDW affinity (**Fig. S15**). We concluded that the shape and volume of this pocket, rather than specific hydrogen bonding interactions with the bound TCR peptide side chain, are important for accommodation of TCR peptide sequences with differing C-terminal amino acids.

**Figure 4.**
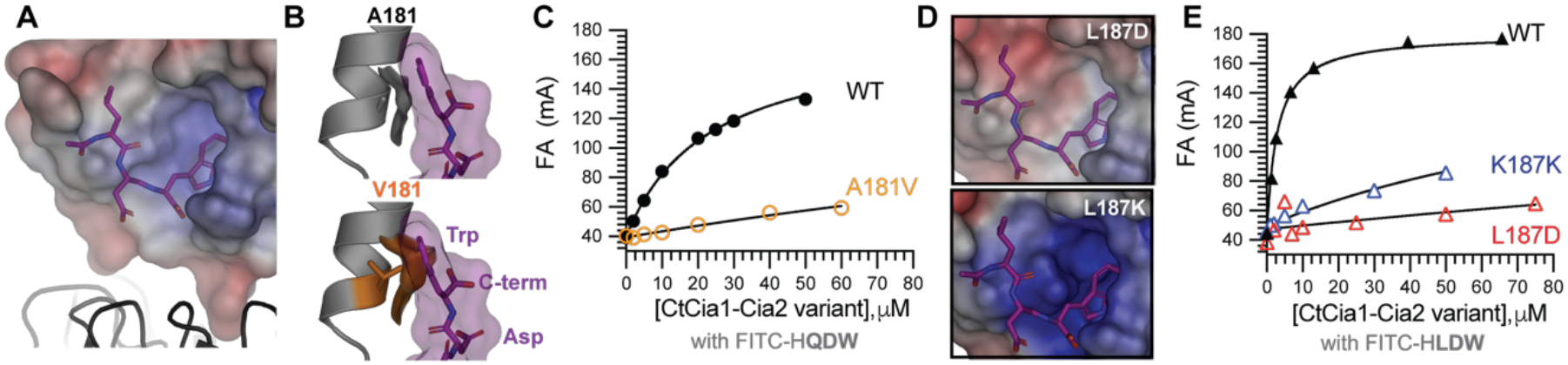
Biochemical validation of TCR peptide binding site. **(A)** View of an MD-derived TCR peptide (purple sticks) binding pose to illustrate electrostatic potential of the *Dm*Cia2 surface surrounding the putative binding site. *Dm*Cia1 is shown as a ribbon. **(B)** Structural comparison illustrating the effect of reducing the volume of the pocket proposed to accommodate the C-terminal aromatic side chain. The surface of the modeled TCR peptide (purple) is shown with Ala181 of *Ct*Cia2 (gray sticks/surface, top) and the Val181 substitution (orange sticks/surface, bottom), which decreases the available pocket volume. **(C)** Fluorescence anisotropy (FA) binding measurements showing reduced affinity of the *Ct*Cia1-Cia2^A181V^ variant for FITC-HQDW probe compared to the wild-type (WT) protein. **(D)** Electrostatic surface in the region of TCR peptide binding upon substitution of *Ct*Cia2 Leu187 with aspartate (top) or lysine (bottom), illustrating how these substitutions alter the electrostatic environment surrounding the TCR peptide. **(E)** FA binding measurements showing reduced affinity of the *Ct*Cia2 Leu187 variants for the FITC-HLDW probe.

The computational modeling also identified a second possible interaction surface, a shallow hydrophobic groove on Cia2 positioned to engage the antepenultimate aliphatic residue of the TCR motif (**Fig. 4A**). In the MD simulations, Asn141 of *Dm*Cia2b (corresponding to Asn184 of *Ct*Cia2) appeared to make polar contacts with the TCR peptide backbone, potentially supporting orientation of the Leu side chain towards the hydrophobic groove (**Fig. 3C**). Thus, we attempted substitute Asn184 of *Ct*Cia2; however, only the serine variant could be isolated suggesting a critical role for this residue for Cia2 structural stability. This *Ct*Cia2-Cia2^N184S^ variant had no measurable impact on TCR peptide binding, likely because the serine side chain retains some hydrogen bonding capacity relative to the native asparagine residue (**Fig. S15C**).

We therefore examined how perturbation of the identified hydrophobic groove impacts TCR peptide binding by substituting Leu187 of *Ct*Cia2 (Leu144 *Dm*Cia2b) with lysine or aspartate (**Fig. 4D**). Both *Ct*Cia2 variants strongly disrupted binding of the FITC-HLDW probe to *Ct*Cia1-Cia2, with aspartate substitution resulting in approximately 200-fold reduction in binding affinity (**Fig. 4E, Fig. S15D-E, Table S6**). Collectively, our biochemical data confirm that the hydrophobic groove identified to contact the -2 aliphatic residue constitutes an essential energetic contributor to TCR peptide binding. Together with perturbations of the aromatic residue binding pocket, these biochemical measurements support the binding site architecture predicted by the computational models.

## Discussion

All Fe-S cluster biogenesis systems face a fundamental challenge: they must selectively recruit dozens of client proteins while maintaining sufficient plasticity to accommodate substantial variation in client sequence and structure. Although short peptide motifs have been implicated in driving Fe– S client recognition in CIA and other Fe–S maturation pathways,(11, 26) the molecular principles by which these signals are decoded have remained poorly defined. Here, we define the physicochemical decoding mechanism by which the CIA targeting complex recognizes the TCR motif. Rather than relying on a fixed consensus sequence, this system interprets a hierarchy of binding determinants to achieve selective yet adaptable client recruitment.

Quantitative energetic dissection of the interaction of TCR peptide with the Cia1-Cia2 complex allowed us to define this decoding grammar. Our biochemical data and MD simulations point to the C-terminal main-chain carboxylate of the TCR motif as a major binding determinant, acting as a non-substitutable anchor making a buried salt bridge with a conserved arginine residue of Cia2 (R134, *Dm*Cia2b).(12) Neutralization of the negative charge or its relocation to a side chain abolishes detectable binding (**Fig. 2A**), underscoring both its essential electrostatic role and strict geometric constraints imposed by acid anchor of the decoding site (**Fig. 5**). This electrostatic contact thereby positions the terminal aromatic side chain of the TCR peptide within a hydrophobic cavity at the Cia1-Cia2 interface. Together, the primary pocket and acid anchor provide the dominant interactions governing TCR peptide recognition (**Fig. 5**). The outsized importance of this minimal recognition element provides molecular context for the earlier observations that mutation or deletion of the C-terminal residue of the CIA clients including viperin, Apd1, and Leu1, or the CIA adaptor Lto1, can disrupt TCR peptide mediated FeS cluster maturation.(11, 27-29)

**Figure 5.**
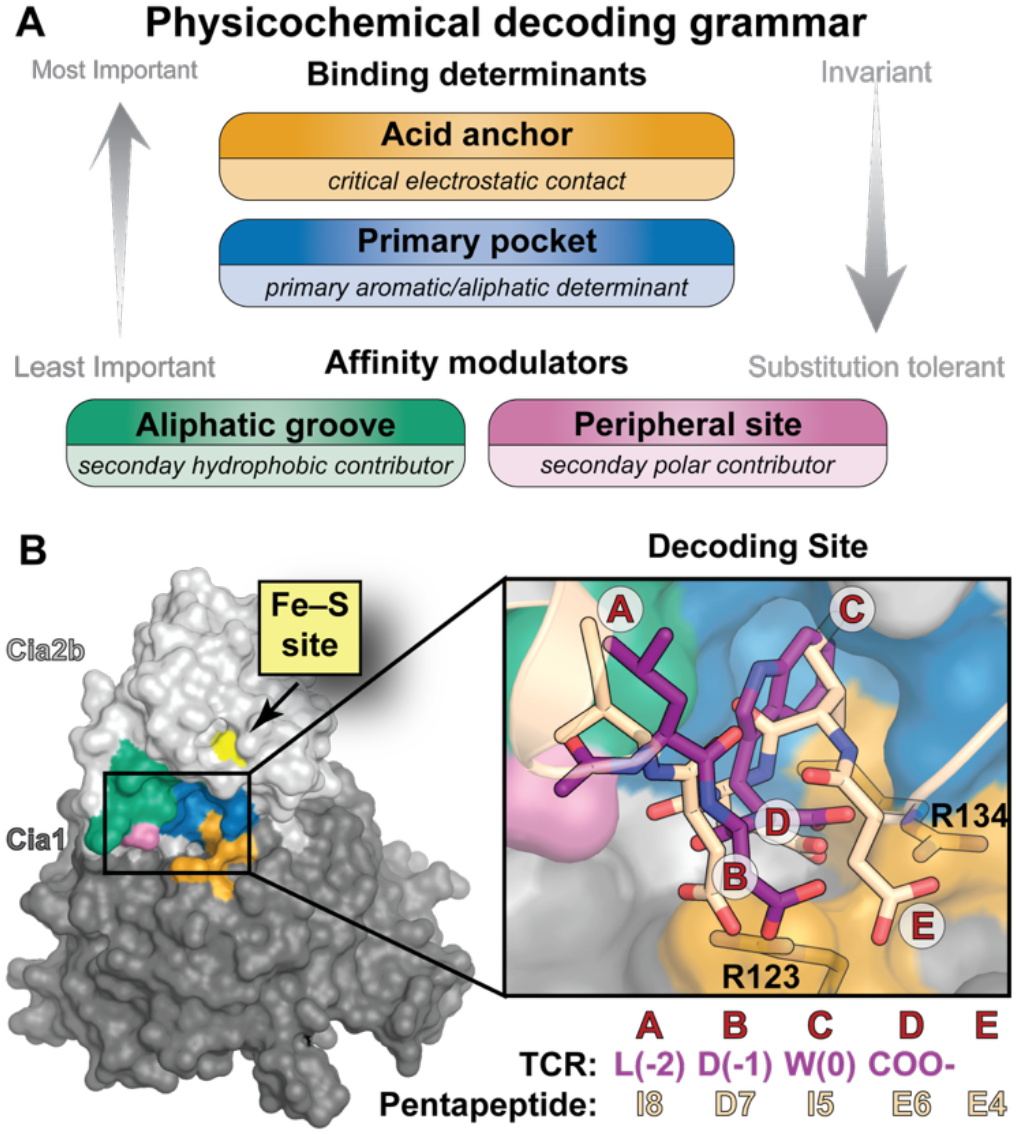
Targeting peptide decoding is governed by physicochemical properties rather than strict sequence constraints. (**A**) Summary of the decoding grammar guiding targeting peptide recognition. Core peptide recognition elements (the acid anchor and primary pocket) provide the dominant binding interactions and are the least tolerant of substitution, whereas the aliphatic groove and peripheral site function as secondary affinity modulators and are more permissive to non-consensus residues. (**B**) The targeting peptide decoding site comprises four regions: the acid anchor (orange), primary pocket (blue), aliphatic groove (green) and the peripheral polar site (pink). The putative Fe–S cluster binding site (yellow) lies adjacent to the decoder, suggesting possible coupling of client binding and cluster delivery. Notably, this physicochemical decoding framework can accommodate other peptides engaging in distinct orientations, as illustrated by the inset comparing TCR peptide binding pose derived from MD simulations (purple sticks) with the proposed internal targeting pentapeptide (wheat sticks). Each moiety of the targeting peptides are labeled A-E to illustrate how chemically similar entities overlap in the two binding poses.

One unexpected outcome of our work is the discovery that the C-terminal residue’s contribution to the binding energy of the TCR peptide is tunable. Although tryptophan and phenylalanine are both consensus residues, they differ substantially in binding affinity. Tryptophan provides maximal affinity, phenylalanine supports intermediate binding affinity, and tyrosine produces weaker but still measurable binding (**Fig. 2C**). These findings align with prior observations that substitution of Leu1’s C-terminal tryptophan with tyrosine reduces but does not abolish TCR-dependent metallocluster maturation.(11) Our expanded bioinformatic analysis further identifies naturally occurring TCR motifs terminating in tyrosine (**Fig. S7**), suggesting that any terminal aromatic residue is compatible with CTC recruitment thus expanding the permissible TCR sequence space.

Residues upstream of the terminal hydrophobic residue further tune binding affinity through sequence context-dependent interactions. The antepenultimate (-2) residue enhances affinity, likely engaging a hydrophobic groove on Cia2 (**Fig. 5**). This additional binding energy can compensate for the weaker contributions from other positions, such as those involving a terminal phenylalanine or tyrosine. This provides the first mechanistic explanation for the enrichment of aliphatic residues at the -2 position of the TCR motif.

In contrast, the -1 position is more tolerant of substitution than sequence conservation alone would suggest. Non-conservative substitutions including introduction of amino acids with large aliphatic side chains or basic residues in place of the penultimate aspartate result in only modest changes in TCR peptide affinity when tryptophan is the terminal residue, but can reduce binding affinity by more than an order of magnitude in Phe-terminated peptides (**Fig. 2D**). MD simulations provide a rationalization for this behavior, revealing fewer electrostatic contacts between the penultimate aspartate of the TCR peptide and Arg123 (*Dm*Cia1 numbering) as compared to the interaction between the main-chain carboxylate and Arg134 (*Dm*Cia2b numbering). Bioinformatic analysis further corroborates this sequence-context dependence. Lysine is enriched at the penultimate position of Trp-terminated TCR sequences but excluded from those ending in Phe. Together these data indicate that Trp- and Phe-terminated targeting peptides engage the CTC via related yet distinct binding modes that allow binding energy to be redistributed across the motif.

Notably, the decoding site lies adjacent to the putative Fe–S cluster binding site of Cia2 (yellow, **Fig. 5A**).(30) This proximity suggests that targeting peptide recognition and Fe–S cluster transfer may be functionally coupled, as previously suggested for the iron-regulated interaction of CTC with Nar1/CIAO3, the putative CIA Fe–S cluster carrier.(31) One important caveat of our work is that the CTC components used herein are in the apo-state as we are unable to produce holo-CTC in a homogenous state.(30, 32) Although our group and others have observed that the CTC can bind clients in the absence of a metallocluster,(10, 12, 33) elucidating how metallocofactor binding influences the affinity and dynamics of the CTC-client interaction is the next important frontier for understanding Fe–S protein maturation.

Structural modeling constrained by biochemical data identified a TCR peptide binding site spanning the Cia1-Cia2 interface. Based on the subsequent MD simulations, the terminal carboxylate is pinned by arginine residues contributed by both subunits, the terminal aromatic side chain nests within a surface cavity on Cia2, and the -2 residue is adjacent to a hydrophobic groove along the same surface (**Fig. 5**). Importantly, pharmacophore mapping using E-FTMap detected strong hydrogen-bond acceptor hotspot consistent with the acid anchor site and a prominent hydrophobic pocket predicted to accommodate both aromatic and aliphatic moieties. The close correspondence between these computationally predicted pharmacophore clusters and the experimentally derived hierarchy reinforces the conclusion that the Cia1-Cia2 interface is structurally organized to decode peptides presenting alternating hydrophobic and acidic features, with the acid anchor and primary pocket providing the dominant interaction sites.

While this manuscript was in preparation, a 2.5 Å crystal structure of the Cia1-Cia2 complex bound to an internal targeting peptide representing an internal targeting motif was reported.(34) Despite its distinct sequence (D-[ILMV]-E-[DEN]-X, where X is any nonpolar or small polar residue) and antiparallel orientation relative to the TCR tripeptide, this internal targeting occupies the same interfacial site identified and engages the same decoding surfaces identified here (**Fig. 5B, Fig S16**). These observations reinforce that targeting peptide recognition is governed by physicochemical features rather than a specific sequence motif.

At the same time, differences between the internal and terminal targeting peptides highlight the constraints imposed by the decoding site. Something here about how steric bulk at the acid anchor appears to shift the alpha carbon of the residue occupying the primary pocket deeper, thus explaining why rigid aromatic groups cannot be substituted for I5. Remarkably, the internal targeting peptide mimics the binding of the Cia2-derived pentapeptide identified herein (**Fig. 3E**). However, SPC-presented variants of this Cia2-derived peptide did not exhibit detectable binding in our assays, indicating that its interaction with the targeting complex is either weak or dependent on structural context not captured by the isolated pentapeptide. Nevertheless, our identification of a Cia2-derived peptide occupying the interfacial binding site supports our model that the Cia1-Cia2 interface functions as a versatile decoding site capable of engaging multiple peptide types provided they share common physiochemical features.

## CONCLUSIONS

Altogether these data support a model in which the Cia1-Cia2 interface functions as a targeting peptide decoder that interprets a hierarchical set of physicochemical features rather than a fixed sequence motif. The acid anchor and hydrophobic pocket define the core recognition elements of this site. Interactions within the hydrophobic groove and peripheral region modulate affinity in a sequence-context-dependent manner, enabling recognition of diverse peptide sequences. Notably, this decoding site accommodates both terminal and internal targeting peptides, despite differences in sequence and binding orientation, highlighting the adaptability of the hierarchical decoding grammar.

At the systems level, recruitment of CIA clients likely relies on multiple points of contact with the targeting complex, not only because a single decoding site is unlikely to provide sufficient specificity to identify dozens of CIA clients in the cellular environment, but also because multiple interaction interfaces are needed to enforce the precise positioning required for Fe–S cluster transfer from the targeting complex to the client. Because other Fe–S cluster pathways also use a single targeting factor to identify many client proteins, the principles established here may shed light on how these systems carry out the final steps of Fe–S protein maturation. More broadly, these findings provide a foundation for understanding how multiprotein assemblies recognize diverse binding partners through hierarchies of physicochemical interactions and degenerate peptide motifs, rather than one-to-one interactions dictating rigid sequence constraints.

## Materials and Methods

### Protein expression, mutagenesis, and purification

The *Chaetomium thermophilum* (*Ct*) Cia1-Cia2 complex and SUMO peptide carrier (SPC) constructs were expressed and purified as described previously, (11, 12) except an avi-tag was introduced at the N-terminus of the SPC (**Detailed Materials and Methods** in **Supporting Information**). For purification of *Saccharomyces cerevisiae* Pri1, the expression clone was created in the pDEST17 vector using a donor plasmid with the *Sc*Pri1 coding sequence in pDONR201 (from the plasmid repository at Arizona State University), transformed into the Rosetta(DE3) strain, and purified via IMAC as described in **Detailed Materials and Methods**. All site-directed mutants were introduced by Q5 mutagenesis (New England Biolabs) and verified by DNA sequencing.

### Fluorescence anisotropy binding assays

Direct fluorescence anisotropy (FA) binding assays were performed using FITC-labeled tetrapeptides (GenScript; 0.05-0.1 µM) with *Ct*Cia1-Cia2 (0-50 µM). The anisotropy values were measured, normalized and fitted as described.(12) For the competitive FA experiments, the FITC-labeled tetrapeptide probe (0.05-0.1µM) was displaced from *Ct*Cia1-Cia2 via an unlabeled competitor, including N-acetyl-amino acids, peptides, or SPC constructs. Concentration of N-acetyl-amino acids was determined using reported extinction coefficients.(35) The anisotropy values at each concentration of competitor were plotted and fitted with a competitive binding model (see **Detailed Materials and Methods**).

### Affinity co-purification assays

Protein-protein interaction analysis via affinity copurification was performed by mixing equimolar amounts of a strep-tagged bait and prey proteins, separation via streptactin chromatography, and the SDS-PAGE analysis with Coomassie staining of input and elution samples as described.(11)

### Bioinformatic analysis

Phe-terminated (n=87) and Trp-terminated (n=88) sequences from CIA client proteins derived from organisms across the eukaryotic domain of life(36) were analyzed separately, and the frequency and enrichment (frequency normalized by frequency in the human proteome) of the amino acids within the TCR motif was determined as described.(11, 12)

### Computational modeling

Detailed description of peptide docking and molecular dynamics simulations are provided in **Detailed Materials and Methods**. Briefly, potential peptide binding hotspots on the *Ct*Cia1-Cia2 interface were identified using the E-FTMap algorithm.(18, 19) Docking of an N-acetyl-LDW tripeptide was performed using the FRED docking program,(20-23) restricting the search region to the interfacial hotspot identified by FTMap. Docked poses were clustered by RMSD and the resulting clusters were visually inspected to identify those consistent with experimentally derived constraints. Molecular dynamics (MD) simulations were performed to evaluate the stability of candidate peptide poses using GROMACS 2018.3, with the topology of a protein–peptide complex was prepared using parameters from the AMBER99SB force field and TIP3P water model.(37-41) LINCS algorithm was used to constrain covalent bonds involving hydrogen atoms.(42, 43)

AlphaFold3 modeling was performed using the available webserver.(17) Figures were generated via Pymol3, with electrostatic surface potential using the APBS electrostatics plugin.(44)

## Acknowledgments

This work was supported by grants R35GM118078 (SV), R43GM144992 (SV), and R01GM121673 (DLP) from the National Institute of General Medical Sciences and by the Boston University Undergraduate Research Opportunities Program (SK and LY).

## References

1. J. J. Braymer, S. A. Freibert, M. Rakwalska-Bange, R. Lill, Mechanistic concepts of iron-sulfur protein biogenesis in Biology. Biochim Biophys Acta Mol Cell Res 1868, 118863 (2021).

2. M. Blahut, E. Sanchez, C. E. Fisher, F. W. Outten, Fe-S cluster biogenesis by the bacterial Suf pathway. Biochim Biophys Acta Mol Cell Res 1867, 118829 (2020).

3. A. Kairis, B. D. Neves, J. Couturier, C. Remacle, N. Rouhier, Iron-sulfur cluster synthesis in plastids by the SUF system: A mechanistic and structural perspective. Biochim Biophys Acta Mol Cell Res 1871, 119797 (2024).

4. V. D. Paul, R. Lill, Biogenesis of cytosolic and nuclear iron-sulfur proteins and their role in genome stability. Biochim. Biophys. Acta 1853, 1528–1539 (2015).

5. M. S. Petronek, B. G. Allen, Maintenance of genome integrity by the late-acting cytoplasmic iron-sulfur assembly (CIA) complex. Front Genet 14, 1152398 (2023).

6. H. Ding, Iron-sulfur cluster biogenesis and regulation of intracellular iron homeostasis in Escherichia coli. Metallomics 17, mfaf040 (2025).

7. N. Maio, T. A. Rouault, Outlining the Complex Pathway of Mammalian Fe-S Cluster Biogenesis. Trends Biochem. Sci 45, 411–426 (2020).

8. K. Gari et al., MMS19 links cytoplasmic iron-sulfur cluster assembly to DNA metabolism. Science 337, 243–245 (2012).

9. O. Stehling et al., MMS19 assembles iron-sulfur proteins required for DNA metabolism and genomic integrity. Science 337, 195–199 (2012).

10. S. A. Kassube, N.H. Thomä, Structural insights into Fe-S protein biogenesis by the CIA targeting complex. Nat. Struct. Mol. Biol. 27, 735–742 (2020).

11. M. D. Marquez et al., Cytosolic iron-sulfur protein assembly system identifies clients by a C-terminal tripeptide. Proc. Natl. Acad. Sci. USA 120, e2311057120 (2023).

12. A. Buzuk et al., A Conserved Cia1-Cia2 Interface Mediates Client Recruitment in the Cytosolic Iron-Sulfur Cluster Assembly Pathway. J. Am. Chem. Soc. 147, 34372–34380 (2025).

13. K. Van Roey et al., Short linear motifs: ubiquitous and functionally diverse protein interaction modules directing cell regulation. Chem Rev 114, 6733–6778 (2014).

14. C. P. Wigington et al., Systematic Discovery of Short Linear Motifs Decodes Calcineurin Phosphatase Signaling. Mol Cell 79, 342-358.e312 (2020).

15. N. E. Davey, L. Simonetti, Y. Ivarsson, The next wave of interactomics: Mapping the SLiM-based interactions of the intrinsically disordered proteome. Curr Opin Struct Biol 80, 102593 (2023).

16. L. Guan, A. E. Keating, Training bias and sequence alignments shape protein-peptide docking by AlphaFold and related methods. Protein Sci. 34, e70331 (2025).

17. J. Abramson et al., Accurate structure prediction of biomolecular interactions with AlphaFold 3. Nature 630, 493–500 (2024).

18. O. Khan et al., E-FTMap: A Protein Structure Based Pharmacophore Identification Server for Guiding Fragment Expansion. J. Mol. Biol. 437, 168956 (2025).

19. O. Khan et al., Expanding FTMap for Fragment-Based Identification of Pharmacophore Regions in Ligand Binding Sites. J Chem Inf Model 64, 2084–2100 (2024).

20. M. McGann, FRED pose prediction and virtual screening accuracy. J Chem Inf Model 51, 578–596 (2011).

21. M. McGann, FRED and HYBRID docking performance on standardized datasets. J. Comput. Aided Mol. Des. 26, 897–906 (2012).

22. B. P. Kelley, S. P. Brown, G. L. Warren, S. W. Muchmore, POSIT: Flexible Shape-Guided Docking For Pose Prediction. J Chem Inf Model 55, 1771–1780 (2015).

23. P. C. Hawkins, A. Nicholls, Conformer generation with OMEGA: learning from the data set and the analysis of failures. J Chem Inf Model 52, 2919–2936 (2012).

24. C. Y. Lee et al., Systematic discovery of protein interaction interfaces using AlphaFold and experimental validation. Mol Syst Biol 20, 75–97 (2024).

25. H. Bret, J. Gao, D. J. Zea, J. Andreani, R. Guerois, From interaction networks to interfaces, scanning intrinsically disordered regions using AlphaFold2. Nat Commun 15, 597 (2024).

26. K. S. Kim, N. Maio, A. Singh, T. A. Rouault, Cytosolic HSC20 integrates de novo iron-sulfur cluster biogenesis with the CIAO1-mediated transfer to recipients. Hum. Mol. Genet. 27, 837–852 (2018).

27. K. Stegmaier et al., Apd1 and Aim32 Are Prototypes of Bishistidinyl-Coordinated Non-Rieske [2Fe-2S] Proteins. J. Am. Chem. Soc. 141, 5753–5765 (2019).

28. A. S. Upadhyay et al., Cellular requirements for iron-sulfur cluster insertion into the antiviral radical SAM protein viperin. J. Biol. Chem. 292, 13879–13889 (2017).

29. V. D. Paul et al., The deca-GX3 proteins Yae1-Lto1 function as adaptors recruiting the ABC protein Rli1 for iron-sulfur cluster insertion. Elife 4, e08231 (2015).

30. V. Maione, F. Cantini, M. Severi, L. Banci, Investigating the role of the human CIA2A-CIAO1 complex in the maturation of aconitase. Biochim. Biophys. Acta Gen. Subj. 1862, 1980–1987 (2018).

31. X. Fan et al., Iron-regulated assembly of the cytosolic iron-sulfur cluster biogenesis machinery. J. Biol. Chem. 298, 102094 (2022).

32. A. T. Vo et al., Defining the domains of Cia2 required for its essential function in vivo and in vitro. Metallomics 9, 1645–1654 (2017).

33. A. Vo et al., Identifying the Protein Interactions of the Cytosolic Iron-Sulfur Cluster Targeting Complex Essential for Its Assembly and Recognition of Apo-Targets. Biochemistry 57, 2349–2358 (2018).

34. W. Ren et al., Client recruitment mechanism of the cytosolic Fe-S cluster assembly targeting complex. EMBO J. 45, 1264–1291 (2026).

35. A. B. Ghisaidoobe, S. J. Chung, Intrinsic tryptophan fluorescence in the detection and analysis of proteins: a focus on Förster resonance energy transfer techniques. Int J Mol Sci 15, 22518–22538 (2014).

36. I. Letunic, P. Bork, Interactive Tree of Life (iTOL) v6: recent updates to the phylogenetic tree display and annotation tool. Nucleic Acids Res. 52, W78–w82 (2024).

37. S. Makeneni, D. F. Thieker, R. J. Woods, Applying Pose Clustering and MD Simulations To Eliminate False Positives in Molecular Docking. J Chem Inf Model 58, 605–614 (2018).

38. S. Páll, M. J. Abraham, C. Kutzner, B. Hess, E. Lindahl (2015) Tackling Exascale Software Challenges in Molecular Dynamics Simulations with GROMACS. In Lecture Notes in Computer Science (Springer International Publishing, Cham), pp 3–27.

39. N. Goga, A. J. Rzepiela, A. H. de Vries, S. J. Marrink, H. J. C. Berendsen, Efficient Algorithms for Langevin and DPD Dynamics. J Chem Theory Comput 8, 3637–3649 (2012).

40. W. L. Jorgensen, J. Chandrasekhar, J. D. Madura, R. W. Impey, M. L. Klein, Comparison of Simple Potential Functions for Simulating Liquid Water. 79, 926–935 (1983).

41. S. A. Showalter, R. Brüschweiler, Validation of molecular dynamics simulations of biomolecules using NMR spin relaxation as benchmarks: Application to the AMBER99SB force field. 3, 961–975 (2007).

42. H. J. C. Berendsen, J. P. M. Postma, W. F. van Gunsteren, A. Di Nola, J. R. Haak, Molecular Dynamics with Coupling to an External Bath. J. Chem. Phys. 81, 3684–3690 (1984).

43. B. Hess, P-LINCS: A parallel linear constraint solver for molecular simulation. J Chem Inf Model 4, 116–122 (2008).

44. E. Jurrus et al., Improvements to the APBS biomolecular solvation software suite. Protein Sci. 27, 112–128 (2018).

