## Supporting Information for "Hierarchical decoding of targeting tripeptide motif by the cytosolic iron-sulfur cluster assembly targeting complex"

*Corresponding author.

Supporting information includes:

Supplemental Text, related to computational modeling of the peptide

Supporting Tables **S1-S6**

Supporting Figures **S1-S15**

Detailed Materials and Methods

### Supplemental Text

#### Computational modeling of N-acetyl Leu-Asp-Trp peptide binding to the Cia1-Cia2 interface

To evaluate the structural basis for TCR peptide recognition, we combined peptide docking using *Drosophila melanogaster* Cia1-Cia2 (PDB: 6TBN) as the receptor model and molecular dynamics (MD) simulations.(10) We focused on two highly ranked E-FTMap hotspot regions at the Cia1-Cia2 interface, an apolar/aromatic pharmacophore cluster and a hydrogen-bond acceptor cluster since these hotspots were reminiscent of the binding determinants we identified experimentally: the C-terminal aromatic side chain and the terminal carboxylate.

Because E-FTMap identifies favorable interaction regions rather than peptide binding poses directly, we next asked whether a TCR peptide could be docked in the region identified by E-FTMap and prior biochemical experimentation.(18, 19) Thus, potential binding poses for the N-acetyl-Leu-Asp-Trp peptide were generated using FRED and clustered by pairwise heavy-atom RMSD using a simple greedy algorithm with a 0.5 Å cutoff. Clusters containing fewer than five members were excluded from further analysis and the remaining 21 clusters were ranked by cluster size and visually inspected to identify those consistent with the following criteria determined from biochemical experiments (**Fig. 3B, Fig. S8**). First, the C-terminal aromatic side chain had to occupy the top-ranked aromatic atomic consensus site predicted by E-FTMap. Second, the terminal carboxylate and/or the aspartate side chain had to lie near the top-ranked hydrogen bond acceptor consensus site predicted by E-FTMap. Since biochemical dissection of TCR peptide binding revealed that neutralization or repositioning of the carboxylate disrupted peptide binding, the second criterion also required that the C-terminal carboxylate would form favorable contacts with a basic residue of Cia1 and/or Cia2. Third, the penultimate aspartate residue was required to remain at least partially solvent exposed, consistent with the modest impact on binding affinity when it was substituted in the Leu-Asp-Trp sequence context. Of the 21 clustered peptide docking poses, only one satisfied all biochemical constraints. This pose was therefore selected as the starting point for subsequent MD simulations.

To examine the conformational behavior of the docked peptide binding pose, we initiated ten independent 100 ns MD simulations from the selected starting structure. Peptide conformations from the final 1 ns of each of these trajectories were extracted and clustered as described for the docked peptide poses, except that 1.0 Å cutoff was used (see **Detailed Materials and Methods**). This analysis identified a small number of closely related peptide conformations and the centroid of the most populated cluster remained consistent with the biochemical constraints. We then performed ten additional 10 ns simulations from the centroid of this most populated initial cluster. Across these trajectories, the peptide remained associated with interfacial region and maintained a relatively stable conformation, with an average peptide RMSD of 1.54 ± 0.6 Å with respect to the starting structure (**Fig. S9**). Representative structures from the most populated clusters generated from this second round of MD simulations showed modest differences in peptide conformations while preserving the overall placement of the C-terminus and the aromatic side chain (**Fig. S9, Fig. S10**). In these representative structures, binding appears to be anchored by the C-terminus and aromatic side chain moieties, whereas the leucine side chain position is more inconsistent. The relative variability of the leucine side chain is consistent with biochemical data indicating that it plays a role in tuning peptide affinity rather than being a primary binding determinant.

#### MD simulations to probe interaction of TCR peptide carboxylates with arginine residues at the Cia1-Cia2 interface

To examine possible electrostatic interactions capable of stabilizing the modeled complex, we next analyzed interatomic distances between peptide carboxylate moieties (C-terminus and aspartate side chain) and arginine residues at the Cia1-Cia2 interface (Arg134 of *Dm*Cia2b and Arg123 of *Dm*Cia1).(12) Since MD simulations placed the C-terminus of the TCR peptide adjacent to Arg134 and E-FTMap analysis identified a hydrogen-bond acceptor hotspot in the same region, we first quantified the distance between these groups in the MD simulations to identify possible polar contacts between these moieties (**Fig. S11**). For the Leu-Asp-Trp peptide, 70.9% of MD frames suggested formation of two polar contacts between the C-terminus and the guanidinium group of Arg134 whereas only 7.8% of MD frames appeared to lack any contact between these moieties (**Fig. S12**), indicating a highly persistent polar interaction.

We also examined the interaction of the penultimate aspartate side chain of the TCR tripeptide. In the initial docked model, the aspartate side chain was positioned near Arg123 (*Dm*Cia1 numbering), consistent with a possible salt bridge. Since our biochemical data indicated that substitution of this residue had only a modest effect on affinity in the Trp-terminated context, we again quantified the distance between the carboxyl group of the peptide aspartate side chain and the guanidinium group of Arg123. In contrast to the interaction of the peptide C-terminus with Arg134 of *Dm*Cia2b which maintained two polar interactions across 70.9% of trajectories, the population of MD frames with interaction of the peptide aspartate with Arg123 were more equally distributed between those consistent with two (28.8%), one (40.7%) or no (30.5%) polar contacts (**Fig. S11**, **Fig. S12**). These quantitative analyses of MD simulation frames indicate that the penultimate aspartate residue of the TCR motif forms a more transient electrostatic interaction with Cia-Cia2 as compared to the peptide C-terminus, consistent with the binding data showing the C-terminus is a critical, invariant binding determinant whereas the aspartate side chain plays a role in tuning affinity and thus is more tolerant of substitution.

### The identified peptide binding mode can accommodate TCR peptide variants

Because biochemical measurements indicated that Phe-terminated motifs can bind at the same site as Trp-terminated motifs, albeit more weakly in otherwise identical sequence contexts, we next examined the effect of substituting the terminal tryptophan with phenylalanine to test whether the same general binding mode could be maintained. Performing 10 independent 10 ns simulations with the starting model of *Dm*Cia1-Cia2 bound the N-acetyl-Leu-Asp-Phe revealed peptide trajectories that remained relatively stable, with an average peptide RMSD of 2.06 ± 0.88 Å relative to the starting structure (**Fig. S9, Fig S10**). Representative structures derived from these simulations indicated that the terminal phenyl side chain remained positioned within the same hydrophobic region occupied by the tryptophan side chain in the simulations with the Trp-terminated peptide. Quantitative analysis of MD frames to monitor interaction of the C-terminal carboxylate with Arg134 of *Dm*Cia2b and the aspartate side chain with Arg123 of *Dm*Cia1 were also similar to that observed with Trp-terminated sequences (**Fig. S11**, **Fig. S12**). We concluded from these simulations that Phe- and Trp-terminated peptides could be accommodated within the same interfacial binding region, consistent with the biochemical analysis.

Finally, we additionally examined peptides with substitutions of the penultimate acidic residue and the -2 leucine. Elimination of the carboxylate side chain by its substitution with alanine did not prevent stable peptide binding at the interfacial binding region (**Fig. S9**), consistent with the biochemical observation that substitution of the penultimate aspartate contributes only modestly to TCR peptide binding affinity. More broadly, comparison of representative structures for antepenultimate Leu to Gln, penultimate Asp to Ala, and terminal Trp to Phe variants indicates that all of these sequence variations can be accommodated at the putative peptide binding site with mainly local changes in the central position of the -2 and -1 residues while the aromatic side chain and C-terminus remained anchored to the same general position occupied by the N-acetyl Leu-Asp-Trp tripeptide at the Cia1-Cia2 interface (**Fig. S9**). In total, our MD simulations support a model in which the peptide C-terminal aromatic residue and its main-chain carboxylate provide the primary binding determinants whereas the penultimate residue is more conformationally flexible and energetically permissive of substitution.

**Table S1. Apparent $K_D$ values for various SPC constructs**

| Unlabeled Competitor | $K_D$, μM [i] | Fold change in affinity [ii] | ΔΔG [ii], kcal/mol |
|---|---|---|---|
| SPC-Leu$1_{21}$ | 32±4.1 (n=2) | - | - |
| SPC-Leu$1_3$ | 17 | ↓0.5 | -0.4 |
| W779F SPC-Leu$1_{21}$ | 510 | ↑16 | 1.6 |
| W779Y SPC-Leu$1_{21}$ | >2500 | ↑>70 | >2.6 |
| W779G SPC-Leu$1_{21}$ | >2500 | ↑>70 | >2.6 |
| W779L SPC-Leu$1_{21}$ | >2500 | ↑>70 | >2.6 |
| SPC-Nar$1_{10}$ | 170 | ↑5.3 | 1.0 |

[i] Apparent $K_D$ values determined by competitive displacement of FITC-HQDW, with the exception of SPC-Leu$1_3$ for which FITC-HLDW was used.
[ii] Relative to SPC-Leu$1_{21}$

**Table S2. Apparent $K_D$ values for N-acetyl amino acids**

| Unlabeled competitor | $K_D$, μM[i] | Fold change in affinity[ii] | ΔΔG[ii], kcal/mol |
|---|---|---|---|
| SPC-Leu$1_{21}$ | 47±5.0 (n=2) | - | - |
| N-acetyl-Trp | 630±3.0 (n=2) | ↓13 | 1.54±0.06 |
| N-acetyl-Phe | ~10,000 | ↓210 | 3.2 |
| N-acetyl-Tyr | >20,000 | ↓>430 | >3.6 |
| N-acetyl-6F-Trp | 2300 (n=1) | ↓49 | 2.3 |

[i] Apparent $K_D$ values determined by competitive displacement of FITC-HQDW from *Ct*Cia1-Cia2
[ii] Relative to SPC-Leu$1_{21}$

**Table S3. Apparent $K_D$ values for various FITC-labeled tetrapapetides**

| FITC Tetrapeptide | $K_D$, μM[i] | Fold change in affinity[ii] | ΔΔG[ii], kcal/mol |
|---|---|---|---|
| HQDW | 23±2.5 | - | - |
| HLDW | 2.5±0.5 (n= 4) | ↑9.2 | -1.31±0.13 |
| HLDF | 50.9±4.3 (n=2) | ↓2.2 | 0.47±0.08 |

[i] Apparent $K_D$ values determined by titrating the indicated FITC-tetrapeptide with *Ct*Cia1-Cia2
[ii] Relative to FITC-HQDW

**Table S4. Apparent $K_D$ values for various SPC-Leu1$_{21}$ TCR tripeptide variants**

| TCR tripeptide sequence [i] | $K_D$, μM [ii] | Fold change in affinity [iii] | ΔΔG[iii], kcal/mol |
|---|---|---|---|
| QDW (WT) | 41±9 (n=6) | - | - |
| **L**DW | 4.0±2.1 (n=4) | ↑10 | 1.38±0.34 |
| QD**F** | 610 | ↓15 | 1.6 |
| **LDF** | 45±4.0 (n=4) | ↓1.1 | 0.1 |
| QD**Y** | >2500 | ↓>60 | >2.4 |
| **L**D**Y** | 130 | ↓3.2 | 0.7 |

[i] SPC-Leu1$_{21}$ with “QDW” motif is the parent (WT) construct
[ii] Apparent $K_D$ values determined by competitive displacement of FITC-HLDW from *Ct*Cia1-Cia2
[iii] Relative to WT SPC-Leu1$_{21}$

**Table S5. Apparent $K_D$ values for TCR tripeptide variants of the penultimate amino acid, corresponding to D778 of Leu1**

| TCR tripeptide sequence [i] | $K_D$, μM [ii] | Fold change in affinity [iii] | ΔΔG[iii], kcal/mol |
|---|---|---|---|
| QDW | 26±7.0 (n=6) | - | - |
| Q**A**W | 75±22 (n=2) | ↓2.9 | 0.63±0.24 |
| Q**V**W | 160 | ↓6.2 | 1.1 |
| Q**L**W | 70 | ↓2.7 | 0.6 |
| Q**K**W | 140±18 (n=2) | ↓5.4 | 1.00±0.18 |
| LDF | 30±2.7(n=4) | - | 0.0 |
| L**A**F | 160±20(n=3) | ↓5.3 | 0.99±0.09 |
| L**S**F | 70 | ↓2.3 | 0.5 |
| L**K**F | 610 | ↓20 | 1.8 |

[i] wild-type (WT) SPC-Leu1$_{21}$ corresponds to the “QDW” tripeptide
[ii] Apparent $K_D$ values determined by competitive displacement of FITC-HLDW from *Ct*Cia1-Cia2
[iii] For Trp-terminated sequences, changes reported are relative to WT SPC-Leu1$_{21}$. For Phe-terminated sequences, changes reported are relative to SPC-Leu1$_{21}$ with a “LDF” tripeptide.

**Table S6. The apparent $K_D$ of *Ct*Cia1-Cia2 variants for various FITC-tetrapeptide probes.**

| *Ct*Cia2 variant | $T_m$, °C | $\Delta T_m$, °C | Tetrapeptide | $K_D$, µM | Fold change [i] | ΔΔG, [i] kcal/mol |
|---|---|---|---|---|---|---|
| WT | 69.5 | - | HQDW | 23 | - | - |
| | | | HLDW | 2.8 | - | - |
| | | | HLDF | 54 | - | - |
| A181V | 68.6 | -0.9 | HQDW | 340 | 15 | 1.6 |
| | | | HLDF | 540 | 10 | 1.4 |
| L187K | 68.9 | -0.6 | HQDW | 230 | 10 | 1.4 |
| | | | HLDW | 75 | 27 | 2.0 |
| | | | HLDF | 990 | 18 | 1.7 |
| L187D | ND[ii] | ND | HLDW | 520 | 190 | 3.1 |
| N184S | 69.0 | -0.5 | | ND | ND | ND |
| T133A | ND | ND | HLDW | 2.0 | 0.71 | |

[i] Compared to the WT *Ct*Cia1-Cia2 for binding the same FITC-tetrapeptide as indicated by the light gray and dark gray shading for HLDW and HLDF tetrapeptides, respectively, or no shading for the HQDW tetrapeptide.

[ii] Not determined, ND

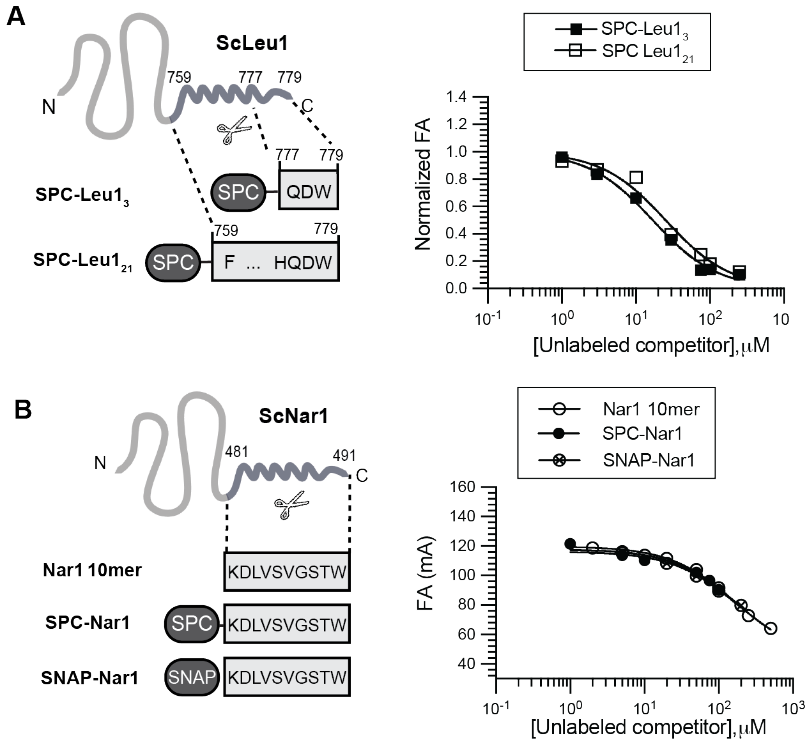


**Figure S1. Validation of the FA assay. (A)** SPC-Leu1 constructs used in the competition FA assay and representative displacement curves, normalized and fitted as described in **Detailed Materials and Methods**. The numbers above the schematic correspond to residue numbering in full-length *Sc*Leu1. The short (SPC-Leu1$_3$) and extended (SPC-Leu1$_{21}$) constructs displaced the FITC-HLDW (0.05 μM) from *Ct*Cia1-Cia2 (3 μM) with apparent $K_D$ values of 17 μM, and 25 μM, for short and extended TCR peptides, respectively, demonstrating that the 18 residues upstream of the TCR tripeptide do not strongly contribute to its binding affinity. **(B)** The SPC carrier display of the TCR peptide does not impact binding affinity. A peptide corresponding to the final ten residues of *Sc*Nar1 was used in the competitive FA assay and compared to the same peptide appended to the SPC or a SNAP tag. All peptides with or without display on a carrier protein displace the FITC-HQDW probe (0.1 μM) from *Ct*Cia1-Cia2 complex (25 μM) with indistinguishable $IC_{50}$ values. The data was fit without normalization and constraining the $A_{min}$ value to 45 mA, the value observed for FITC-HQDW probe in the absence of *Ct*Cia1-Cia2.

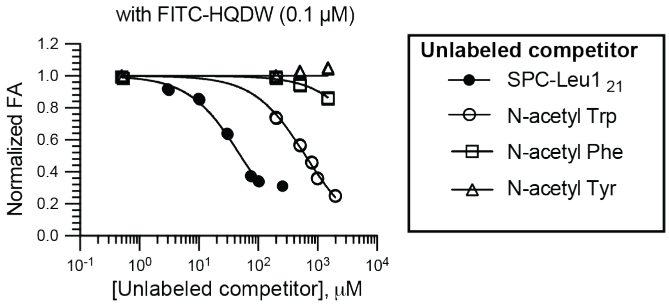


**Figure S2. The C-terminal aromatic residue is a key binding determinant.** Competitive FA assay curves using N-acetylated amino acids as the unlabeled competitor, displacing the FITC-HQDW probe (0.1 µM) from *Ct*Cia1-Cia2 (25 µM). The FA data was normalized and fitted**,** constraining the minimum anisotropy ($A_{min}$) to the value observed for the free (unbound) FITC-HQDW probe (45 mA). In some cases, the data at a high concentration of competitor were excluded from the fit due aggregation that interfered with accurate determination of anisotropy values. The observed $K_D$ values are summarized in **Table S2**.

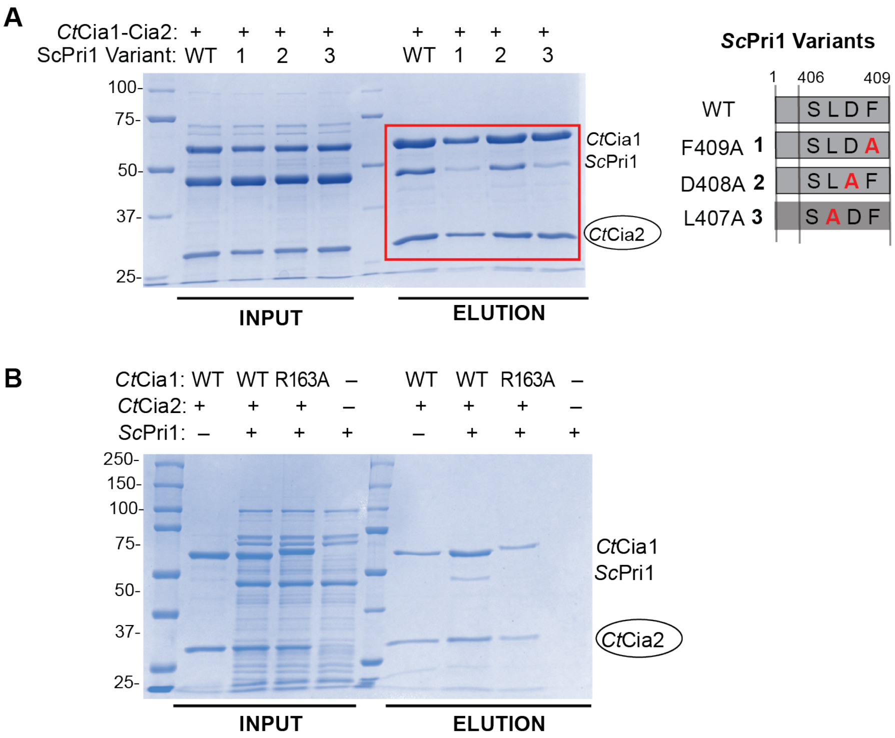


**Figure S3. Pri1's interaction with Cia1-Cia2.** Affinity co-purification analysis of the interaction of *Sc*Pri1 with *Ct*Cia1-Cia2. Wild-type (WT) *Sc*Pri1 or the indicated TCR motif variant (**A**) were mixed with WT *Ct*Cia1-Cia2 or the *Ct*Cia1$^{R163A}$-Cia2 variant (**B**), in the presence (+) or absence (–, no bait negative control) of strep-tagged *Ct*Cia2 (bait, circled). Complexes were isolated using streptactin resin and the input and elution samples were analyzed by SDS-PAGE followed by Coomassie staining. Molecular weight standards (kDa) are indicated to the left. Data shown in **Fig. 2B** is boxed in red.

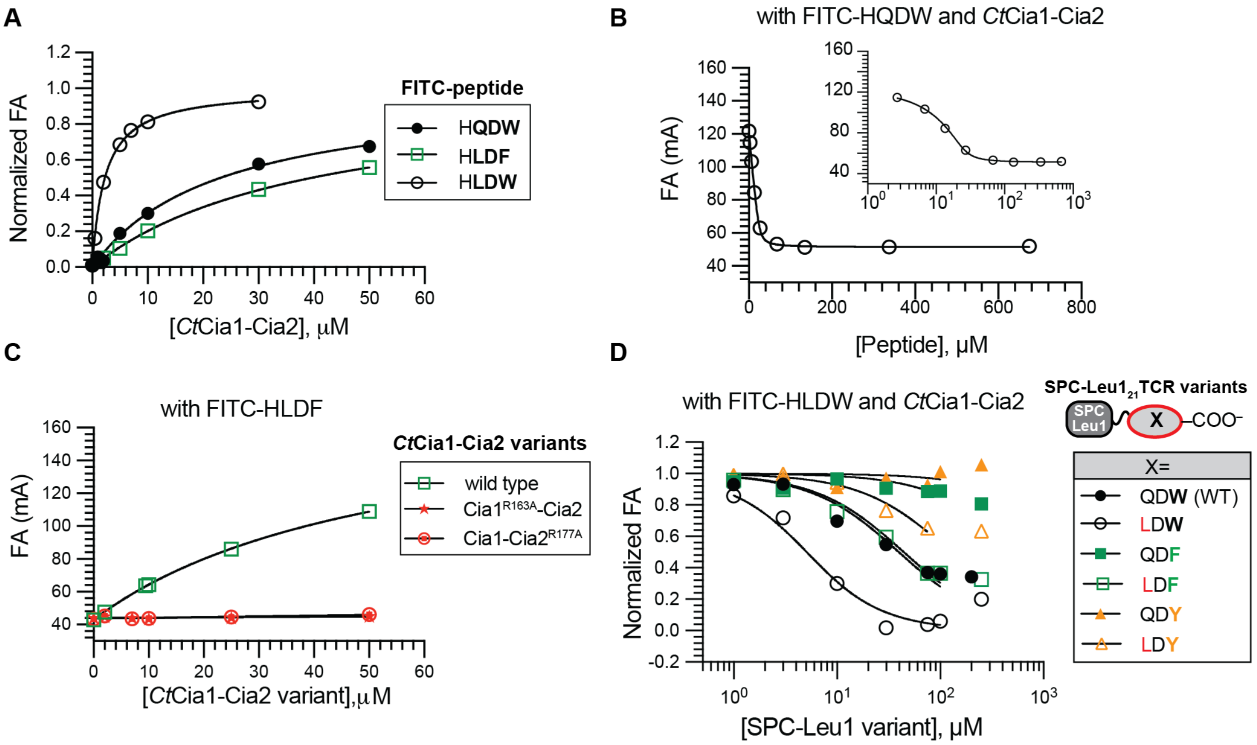


**Figure S4. The antepenultimate (-2) residue of the TCR motif modulates binding affinity. (A)** Fluorescence anisotropy (FA) binding curves for FITC-labeled tetrapeptides. The indicated FITC-peptide (0.05 or 0.1 μM) was titrated with wild-type (WT) *Ct*Cia1-Cia2. The resulting FA values were normalized and fit as to a hyperbolic binding model. The resulting $K_D$ values are reported in **Table S3**. **(B)** Competitive FA assay using the 9mer peptide (TTFDKVH**LDW**) as the unlabeled competitor. The FA data was plotted and fitted to a quadratic binding model, constraining $A_{min}$ to the value observed for the free FITC-HQDW probe (45 mA), yielding a $K_D$ value of 5.3 μM (n=1). The inset shows the same data and fitting, but on a logarithmic scale. **(C)** Binding of the FITC-HLDF (0.1 μM) probe to WT *Ct*Cia1-Cia2 or the indicated variant. The FA data was analyzed as in Panel A. **(D)** Competitive FA assay using the indicated SPC-Leu$1_{21}$ TCR variants. The data was normalized, constraining $A_{min}$ to the value observed when SPC-Leu$1_{21}$ ending in LDW was used as the unlabeled competitor (54 mA). The observed $K_D$ values are summarized in **Table S4**.

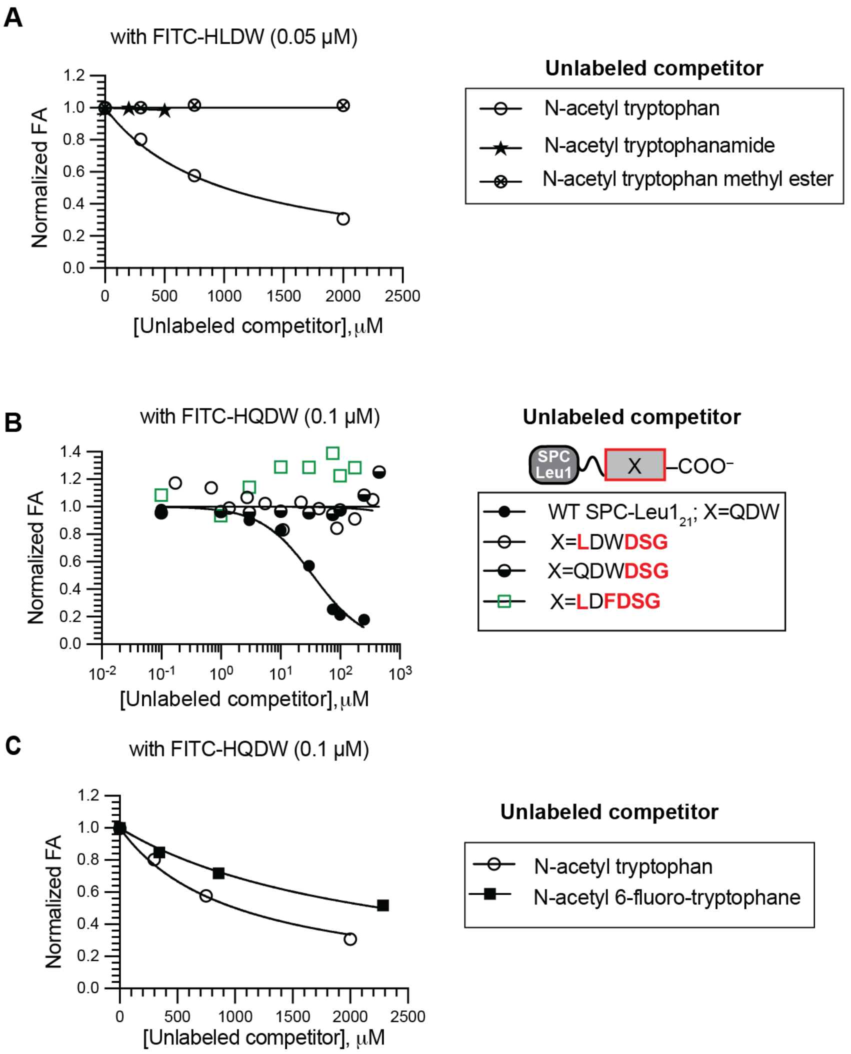


**Figure S5. The main-chain carboxylate serves as a key binding determinant. (A)** Competitive FA curves using N-acetylated Trp or its carboxamide (N-acetyl-L-tryptophanamide) or carboxymethyl ester (N-acetyl tryptophan methyl ester) derivatives as the unlabeled competitor to displace the FITC-HLDW (0.05 μM) probe from *Ct*Cia1-Cia2. The data was normalized and fitted as described in **Materials** and **Methods**. **(B)** Relocation of the carboxylate from the main-chain to a side chain through the addition of a "DSG" tripeptide to the C-terminus of the TCR motif does not recover binding. The indicated SPC-Leu1$_{21}$ variant was used as the unlabeled competitor, displacing FITC-tetrapeptide (HQDW (0.1 μM) for WT and QDWDSG, HLDW (0.05 μM) for LDWDSG and LDFDSG) from *Ct*Cia1-Cia2. The data was normalized constraining $A_{min}$ to the value observed with WT SPC-Leu1$_{21}$ (58 mA) and fitted as described in Materials and Methods. (**C)** Competition FA curves using N-acetylated Trp and its 6-fluoro derivative as the unlabeled competitor displaced the FITC-HQDW (0.1 μM) probe from *Ct*Cia1-Cia2. The data was normalized, constraining $A_{min}$ to the value observed with free FITC-probe (45 mA). Observed dissociation constants were obtained by fitting to a competitive displacement model and are summarized in **Table S2**.

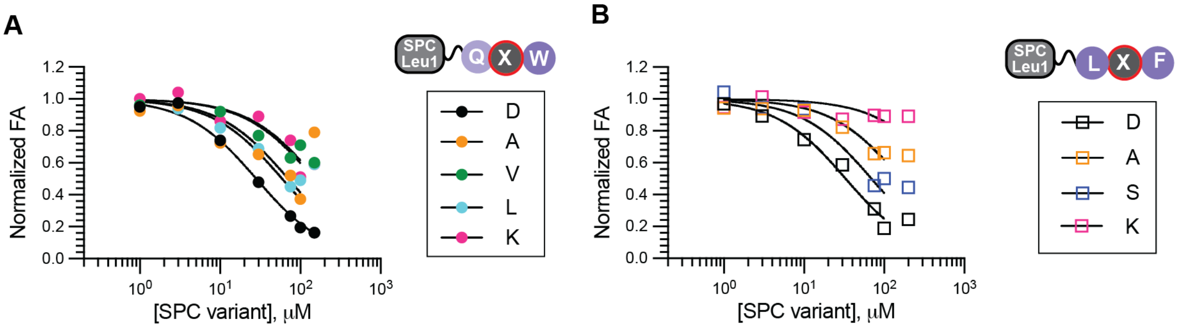


**Figure S6. Penultimate charged residue exhibits high tolerance to substitutions in Trp-terminated TCR sequences.** Competitive FA curves displacing FITC-HLDW (0.05 μM) from *Ct*Cia1-Cia2 with the indicated SPC-Leu1 variants of "QDW" (**A**) and "LDF" (**B**) TCR tripeptide. Cartoon representations of constructs and the legend are located at the right side of each plot. The data was normalized where $A_{min}$ was constrained value observed for wild-type SPC-Leu1$_{21}$ (ending with a QDW tripeptide; 65 mA). The highest concentration points for SPC-TCR constructs were excluded from the fitting due to the observed artifactual increase in FA due to protein instability.

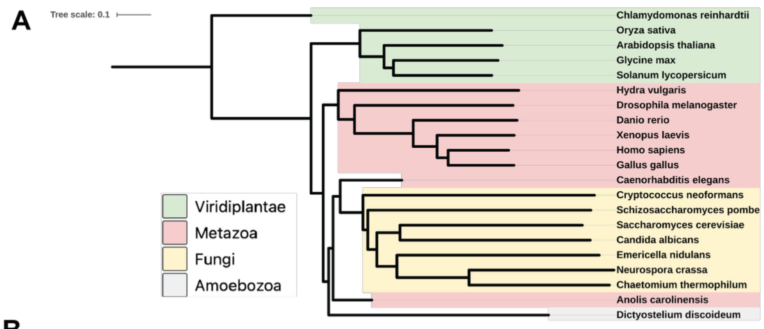


**B**

SPECIES

| Protein NAME Human/Yeast | *Hsa* | *Sce* | *Spo* | *Ani* | *Ncr* | *Dre* | *Xla* | *Cel* | *Dme* | *Ath* | *Gga* | *Cthr* | *Cal* | *Gmx* | *Osa* | *Acs* | *Ddi* | *Cng* | *Cre* | *Sly* | *Hmg* |
|---|---|---|---|---|---|---|---|---|---|---|---|---|---|---|---|---|---|---|---|---|---|
| POLD1/Pol3 | EAW | SKW | MSW | GAW | AMW | FGW | PAW | | AEW | WSF | | AIW | TNW | WNF | WDF | PTW | VEW | TGW | TSW | WNF | FTW |
| REV3L/Rev3 | DQF | NDW | LDW | LEW | LEW | | DNF | KMF | HHF | EWF | DQF | | LDW | EWF | ELF | DQF | FDY | FSW | | | SEF |
| PUR1/Ade4 | LEW | | | | | LEW | LEW | | LSW | | LEW | | | | | LEW | LDF | | | | LQW |
| POLR3F/Rpc6 | LEF | | | LSF | | LDF | LDF | | | LDF | LEF | | | LDF | MDF | LQF | LDF | | LEF | IEF | |
| -/Leu1 | | QDW | NDW | LEW | LEW | | | | | | | LEW | QEW | | | | | TDW | | | |
| RSAD2/- | LDW | | | | | LEW | LEW | | | | LDW | | | | | LEW | | | | | |
| None/Apd1 | | QSW | | ISW | LSW | | | | | | | LSW | IEW | | | | | | | | |
| CTU1/Ncs6 | PTF | LSF | | | | VDF | | LQF | NDF | LDF | | | LSF | LDF | FDF | IFD | | | | | |
| -/- (Neverland) | | | | | | LDF | LDW | FDW | IGW | | | | | | | LDW | | | | | |
| -/- (Nitrite reductase) | | | | IDW | LEW | | | | | | | PDW | | | | | | | | | |
| CIAO3/Nar1 | IRW | STW | NKW | GGW | GGW | IKW | IKW | LKW | INW | NNW | IKW | GGW | SKW | HNW | QNW | IKW | IKW | VVW | ANW | QNW | IQW |
| LTO1/Lto1 | LSF | QSW | | | | MSF | LSF | | YSF | LDF | LTF | | | IQF | | LTF | | | LTF | SGF | |
| PRIM1/Pri1 | KDF | LDF | LEF | MDF | MEF | KDF | KEF | | LQF | FSW | GEF | LEF | LSF | LGW | LTW | MDF | MEF | LDF | | ISW | IDW |
| ELP4/Elp4 | LDF | LDY | LDF | LEF | LDF | LHF | LDF | IDF | LDF | LDF | LDF | | IDF | LDF | LDF | LDF | LDF | YEF | LDF | LDF | LDF |

None | Non Fe-S | Plastidial | Non WFY

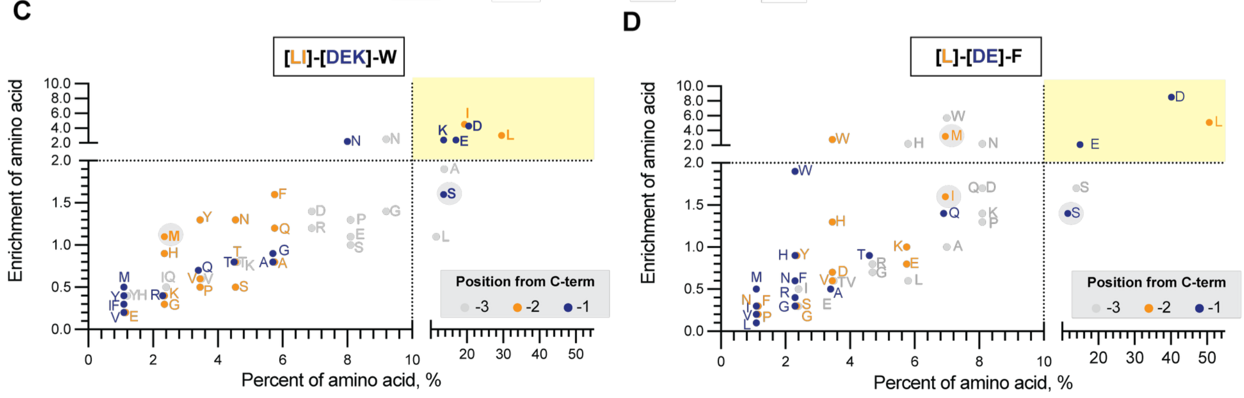


**Figure S7. Bioinformatic analysis of TCR peptides. (A)** Phylogenetic tree of 21 representative eukaryotic species based on Cia1 sequences and generated using by iTol.(32) **(B)** TCR sequences (n=699) of CIA clients and CTC interacting proteins from 8 metazoa (*H. sapiens*, Has; *D. rerio*, Dre; *X. laevis*, Xla; *C. elegans*, Cel; *G. gallus*, Gga, *D. melanogaster*, Dme; *A. carolinensis*, Acs; *H. vulgaris*, Hmg), 5 viridiplantae (*A. thaliana*, Ath; *G. max*, Gmx; *O. sativa*, Osa; *C. reinhardtii*, Cre; *S. lycopercicum*, Sly), 7 fungal (*S. cerevisiae*, Sce; *S. pombe*, Spo; *A. nidulans*, Ani; *N. crassa*, Ncr; *C. thermophilum*, Cthr; C. albican, Cal) and one amoebozoa (*D. discoideum*, Ddi). Cells are color-coded to indicate those that are not Fe–S proteins (yellow), are not not a CIA clients (brown), lack a C-terminal Trp, Phe, or Tyr (dark gray), or for which no homolog identified (light gray). Motifs ending in Tyr are boxed. (**C-D**) Enrichment versus frequency plots for amino acids found at the -1 (blue), -2 (orange), and -3 (gray) position of TCR motifs ending in Trp (**C**, n=88) or Phe (**D**, n=87). Enrichment is calculated relative to the frequency in the human proteome.(11) Residues that are ≥2-fold enriched and found in more than 10% of sequences (upper right quadrant, yellow) are considered consensus. Lysine is enriched at the -1 position in Trp-terminated motifs (13.6%, 2.4-fold enrichment) but excluded from Phe-terminated TCR sequences. Residues previously identified as TCR consensus residues but not meeting these criteria are circled in gray.

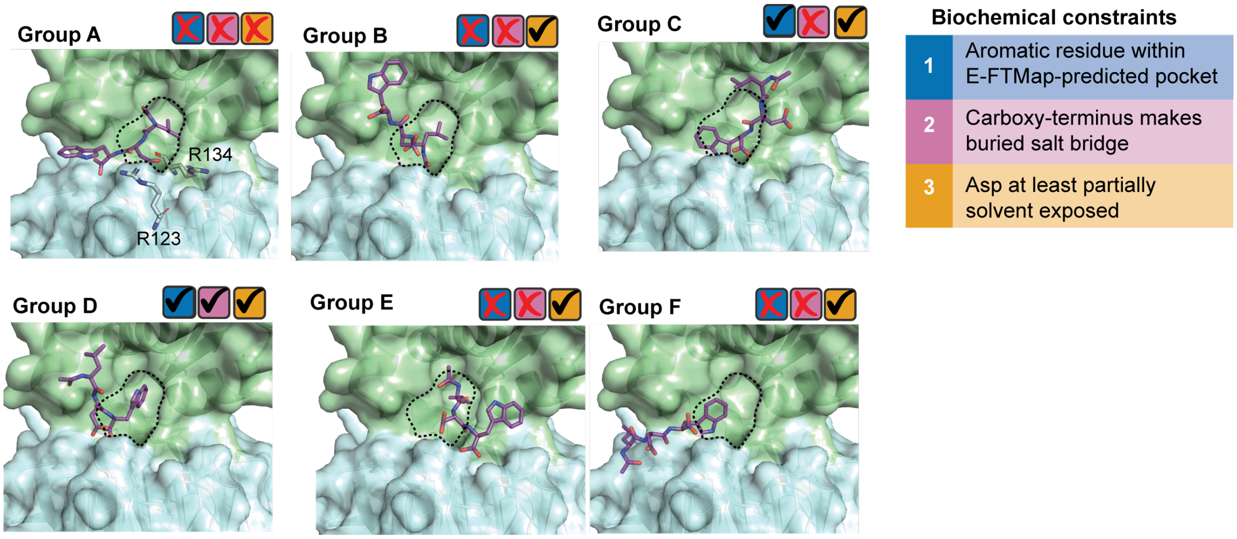


**Figure S8. N-acetylated Leu-Asp-Trp peptide (purple sticks) docked onto Cia1-Cia2** (PDB: 6TBN; *Dm*Cia1, cyan; *Dm*Cia2b, green).(10) Docked peptides were grouped (Group A-F) using a greedy clustering algorithm on the basis of ligand heavy atom RMSD ≤0.5 Å. Possible binding conformations were evaluated for the following criteria: (i) The aromatic side chain binds in the cavity identified by E-FTMap (dashed line, blue checkbox); (ii) C-terminal main chain carboxylate is buried and makes favorable electrostatic interactions (pink checkbox); (iii) the penultimate residue (Asp) is at least partially solvent exposed (orange checkbox). Of the calculated and clustered binding modes, only Group D met all three criteria.

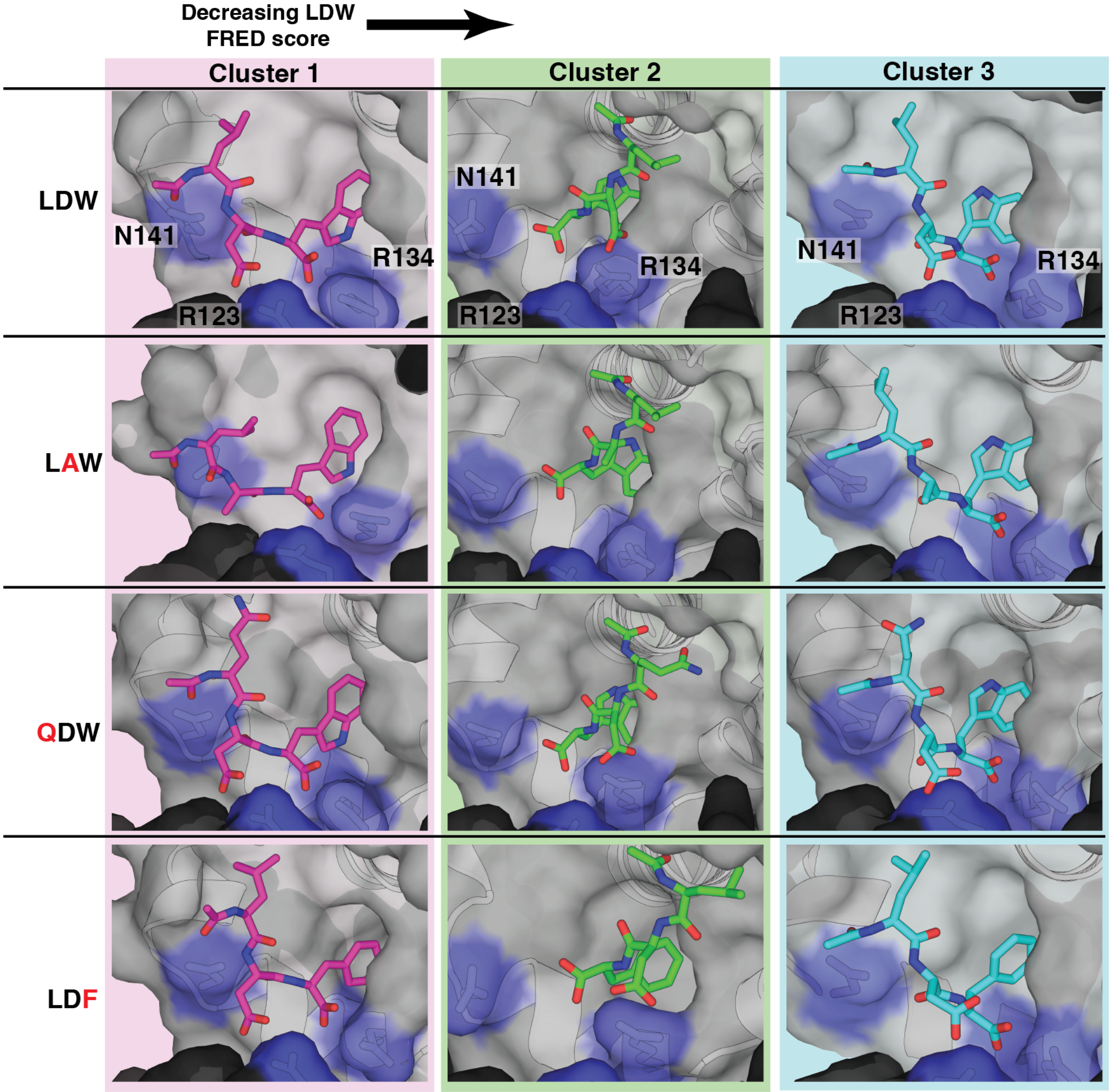


**Figure S9. Representative structures of the most populated MD trajectories (clusters 1-3) for different TCR peptides.** View of an MD-derived TCR peptide (LDW, LAW, QDW, and LDF shown as sticks) binding pose to illustrate the adaptability of the TCR binding site and the preservation of the common binding mode across TCR variants. The surface of residues (*Dm*Cia1, R123 and *Dm*Cia2: R134 and N141) surrounding the binding site are colored in black and purple, respectively.

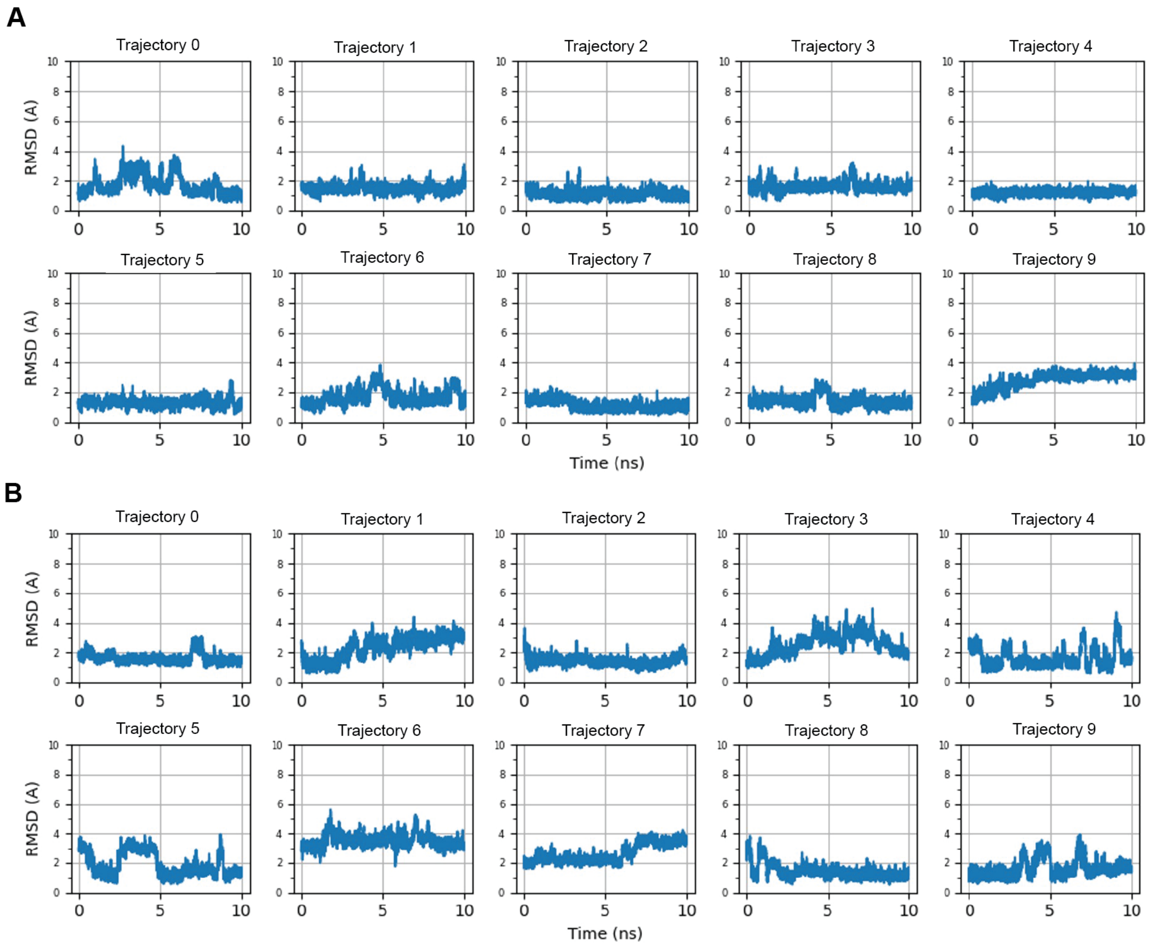


**Figure S10. Root-mean square deviation (RMSD) values for molecular dynamics trajectories.** RMSD values over time the LDW (**A**), and LDF (**B**) tripeptides over the course of MD simulations, calculated relative to the initial docked pose obtained using FRED.

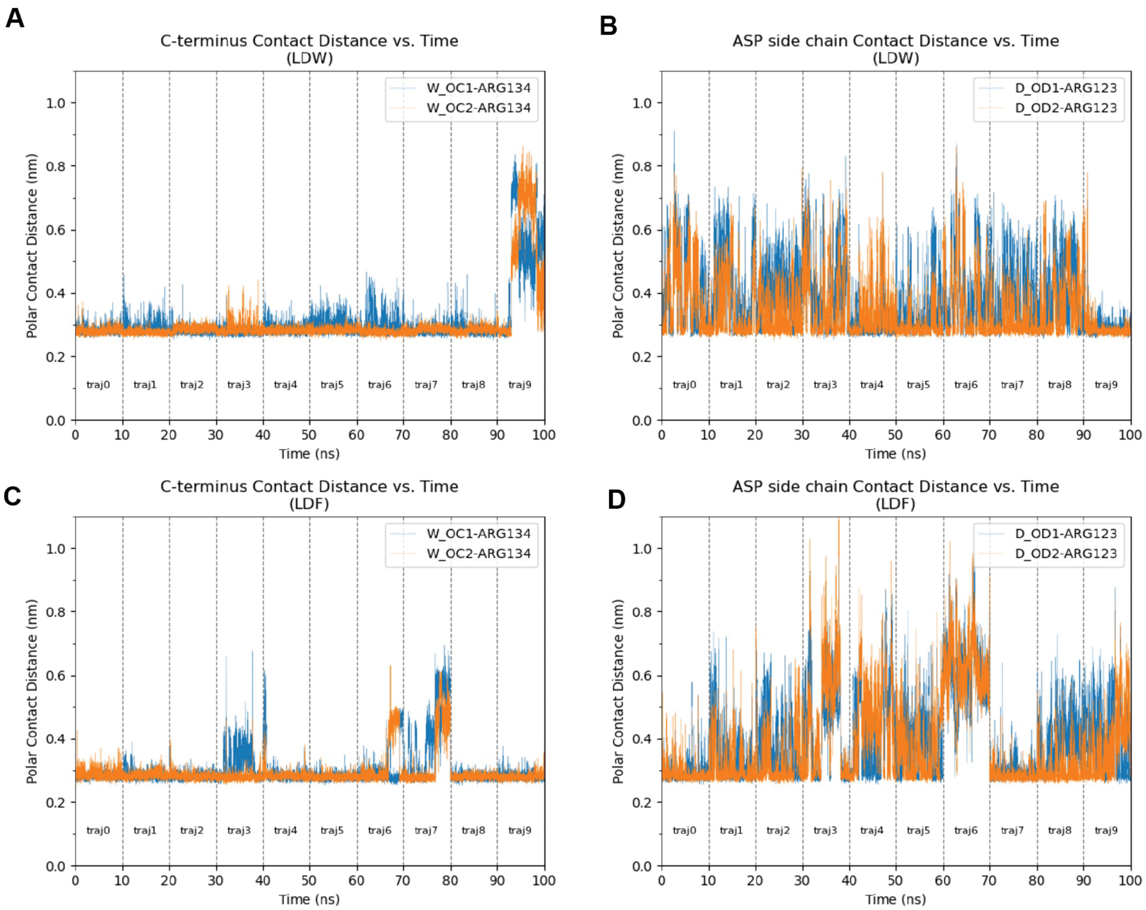


**Figure S11. Minimum distance between oxygen atoms of the carboxy group and nitrogen atoms of the arginine guanidinium group.** Contact distances are plotted over 10 independent 10 ns trajectories for the LDW C-terminal carboxy group with Arg134 of *Dm*Cia2b (**A**), LDW Asp carboxy group with Arg123 of *Dm*Cia1 (**B**), the LDF C-terminal carboxy group with Arg134 of *Dm*Cia2b (**C**), LDF Asp carboxy group with Arg123 of *Dm*Cia1 (**D**). Simulation time is from 10 concatenated molecular dynamics trajectories. W_OC1-ARG134 (blue; panels A and C) and W_OC2-Arg134 (orange, panels A and C) denote distance between each respective carboxy group oxygen and any side chain nitrogen of Arg134 of *Dm*Cia2b. Similarly, D_OD1-ARG123 (blue; panels B and D) and D_OD2-Arg123 (orange, panels B and D) denote distance between each respective carboxy group oxygen and any side chain nitrogen of Arg123 of *Dm*Cia1.

| Moiety | | Percentage of MD frames | | |
|---|---|---|---|---|
| *Guanidinium* | *Carboxylate* | 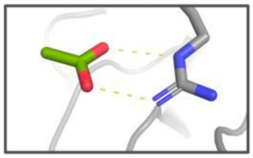 *Two Polar Contacts* | 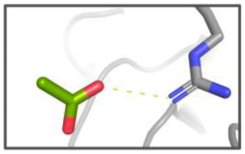 *One Polar Contact* | 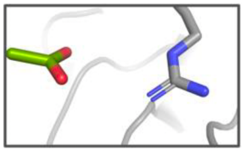 *No Polar Contacts* |
| R134 of *Dm*Cia2 | C-term LDW peptide | 70.9% | 21.3% | 7.8% |
| R123 of *Dm*Cia1 | Asp side chain of LDW peptide | 28.8% | 40.7% | 30.5% |
| R134 of *Dm*Cia2 | C-term LDF peptide | 72.4% | 23.4% | 4.2% |
| R134 of *Dm*Cia2 | Asp side chain of LDF peptide | 24.6% | 40.8% | 34.6% |

**Figure S12. Analysis of polar contacts between TCR carboxylates and CTC arginine residues.** The percentage of MD frames detecting none, one or two polar contacts (as judged by distances as reported in **Figure S11**) between the TCR peptide carboxyl moieties and CTC arginine residues, Arg134 (*Dm*Cia2b) or Arg123 (*Dm*Cia1) as indicated.

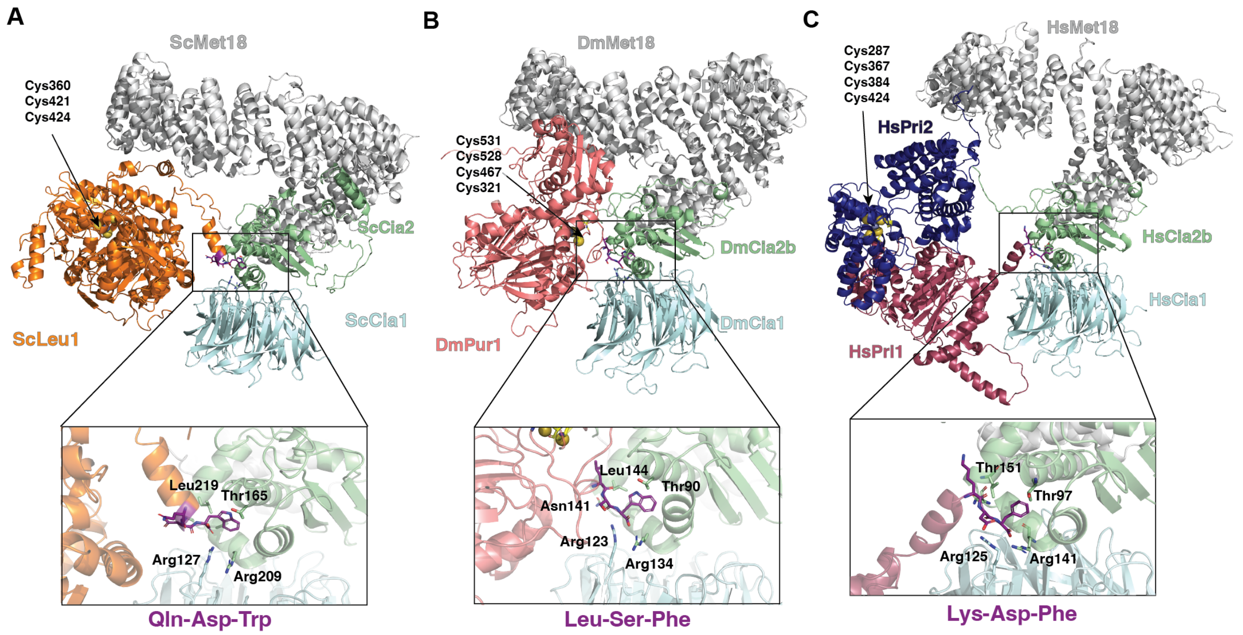


**Figure S13. AlphaFold models CTC bound to various TCR peptide bearing clients.** AlphaFold3-predicted models of the Cia1-Cia2-Met18 complexes from *S. cerevisiae* **(A),** *D. melanogaster* **(B)** and *H. sapiens* **(C)** are shown with the indicated representative CIA client bearing a TCR motif. Each protein ribbon is colored as follows: Cia1, cyan; Cia2, green, Met18, gray, *Sc*Leu1, orange; *Dm*Pur1 salmon; *Hs*Pri2, dark blue, *Hs*Pri1 dark pink. The TCR tripeptide is shown in purple sticks with its sequence indicated in purple text below each inset. Side chains positioned near the TCR motif are shown as sticks colored according to the ribbon color and labeled. Location and numbering of Fe–S binding cysteine residues of CIA clients are indicated.

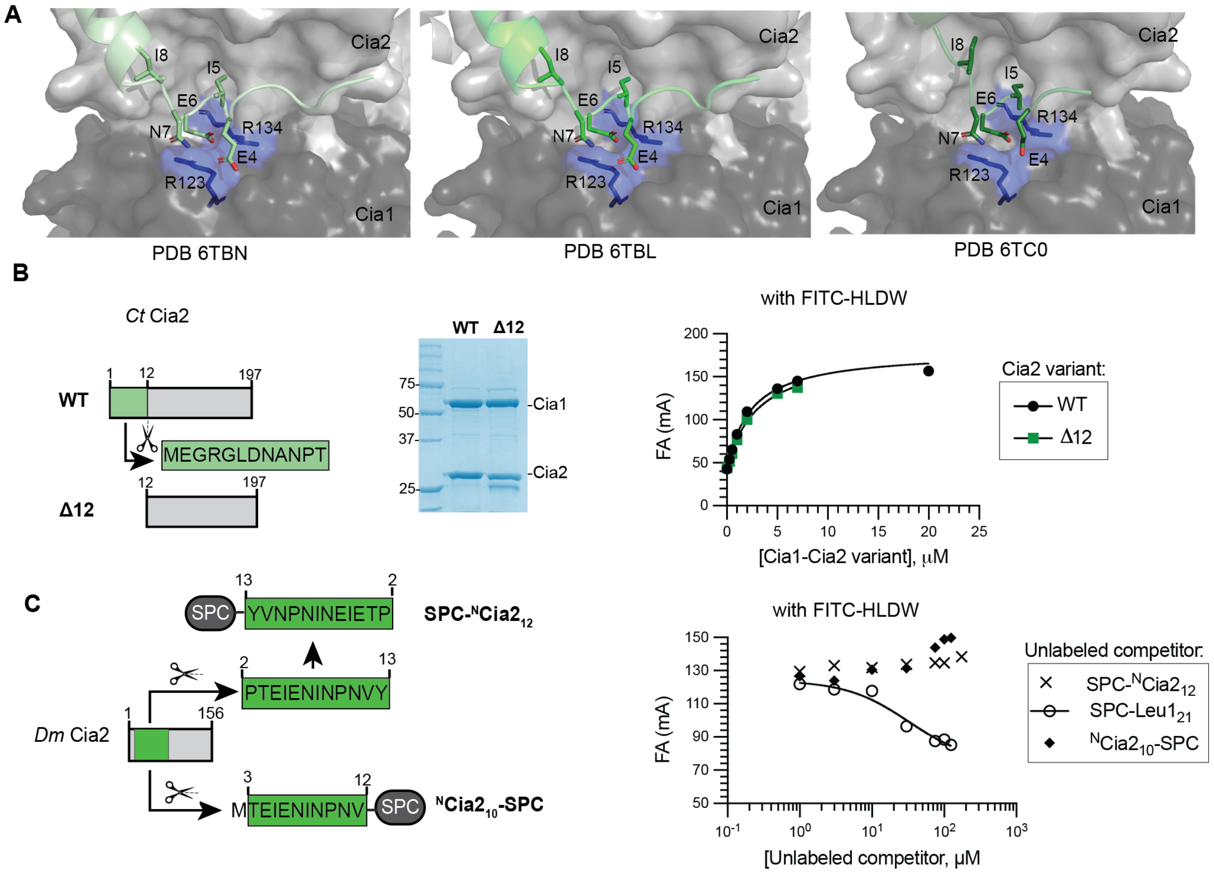


**Figure S14. The N-terminal peptide of DnCia2b occupies the TCR peptide binding site but does not measurably bind *Ct*Cia1-Cia2. (A)** Structural comparison of three *Dm*Cia1-Cia2b crystal structures (PDB: 6TBN, 6TBL, 6TC0) available crystal structures shows a peptide derived from the N-terminus of Cia2b occupies the interfacial TCR peptide binding site. In 6TC0, this peptide originates from one Cia2b protomer and engages the TCR binding site of the opposing protomer within the CTC dimer. In 6TBL and 6TBN, the peptide is contributed by a neighboring crystallographic symmetry mate. **(B)** Deletion of the N-terminal 12 residues of *Ct*Cia2 ($^{\Delta12}$Cia2) does not affect binding of the FITC-HLDW peptide probe to the *Ct*Cia1-$^{\Delta12}$Cia2 complex. Left shows schematic of WT and Δ12 *Ct*Cia2 constructs. Middle shows SDS-PAGE analysis of WT *Ct*Cia1-Cia2 and the Cia1-$^{\Delta12}$Cia2 complex. Right is the FA binding curves showing the WT and Δ12 constructs have similar TCR peptide binding affinities. **(C)** Peptides derived from the N-terminus of *Dm*Cia2b do not compete with TCR probe for binding to *Ct*Cai1-Cia2. Left is schematics of the two SPC constructs to display the EIENI pentapeptide in different orientations. Right is the competitive FA data demonstrating neither SPC construct is competent to competitively displace FITC-HLDW from *Ct*Cia1-Cia2.

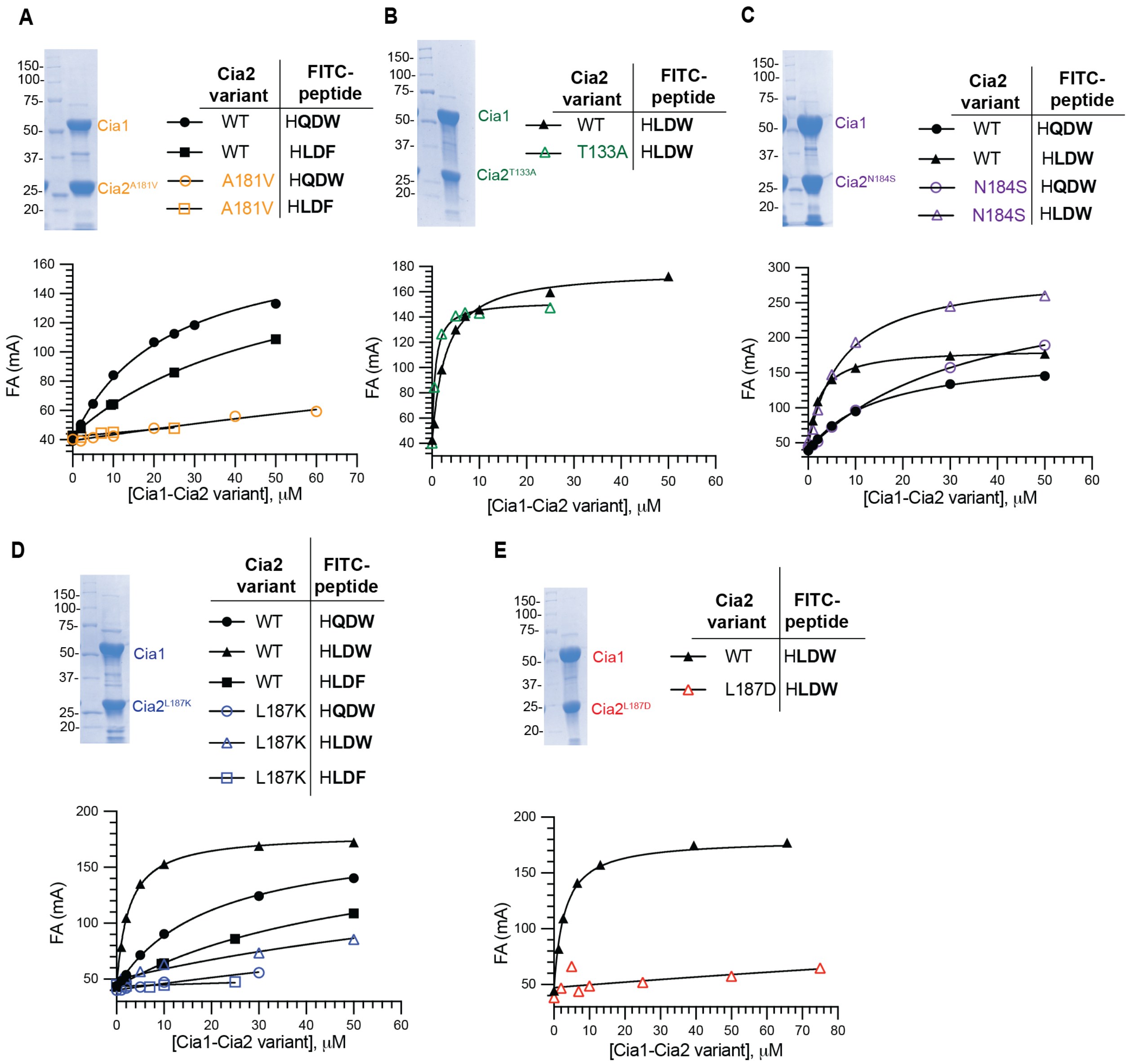


**Figure S15. FA binding assays for *Ct*Cia2 variants.** The binding of indicated FITC-tetrapeptide (● for FITC-HQDW, ▲ for FITC-HLDW, ■ for FITC-HLDF) to WT *Ct*Cia1- Cia2 (in black) or the indicated variant: Cia1-Cia2$^{A181V}$ (orange; **A**), Cia1-Cia2$^{T133A}$ (green; **B**), Cia1-Cia2$^{N184S}$ (purple; **C**), Cia1-Cia2$^{L187K}$ (blue; **D**) and Cia1-Cia2$^{L187D}$ (red; **E**). With the exception of Panel B, data was fit to hyperbolic binding equation, constraining the $A_{max}$ value to observed with the WT Cia1-Cia2 complex (180 mA). For Panel B, no constraints in fitting was applied due to the increased maximum FA observed for this variant, possibly due to nonspecific binding or protein aggregation. The determined $K_D$ values are summarized in **Table S6**.

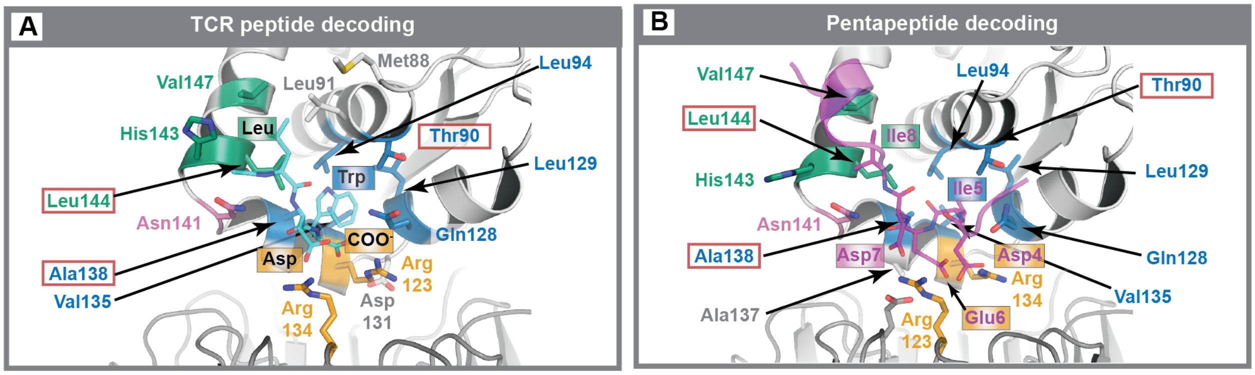


**Figure S16. Structural comparison of terminal TCR and internal targeting pentapeptide binding modes.** Residues within 4 Å of the terminal TCR peptide (**A**, cyan sticks) or internal pentapeptide (**B**, magenta) are shown at the *Dm*Cia1-Cia2b interface. Targeting peptide residues are labeled colored according to their positions within the decoding site as follows: aliphatic groove, green; primary pocket, blue; acid anchor, orange; and peripheral site, pink. Residues from *Dm*Cia1 (dark gray, bottom) or *Dm*Cia2b (light gray, top) within 4 Å of both terminal and internal targeting peptides are also colored according to the subsite they contribute to. Residues found in only one targeting peptide binding site are gray.

### Detailed Materials and Methods

***Ct*Cia1-Cia2 mutagenesis and purification.** The *Ct*Cia2 variants, A181V, N184A/S, T133A, L187D, and L187K, were created by Q5 Site-Directed Mutagenesis Kit (New England Biolabs), using the pETDuet plasmid for co-expression of His-tagged *Ct*Cia1 and Strep-tagged *Ct*Cia2 as the template.(12) All sequences were confirmed by DNA sequencing. Purification and the thermal stability of *Ct*Cia1-Cia2 variants protein were carried out as described.(12)

**SPC cloning and purification.** The plasmid for expression of SPC (SUMO-fused to an N-terminal 6xHis-tag) and C-terminal peptide corresponding to the last 21 amino acids of *S. cerevisiae* (*Sc*) Leu1 (SPC-$Leu1_{21}$) was previously described.(11) An avi-tag (GLNDIFEAQKIEWHE) coding sequence was inserted before the His-tag of the SPC-Leu1 expression vector. Variants were then introduced via Q5 Site-Directed Mutagenesis Kit (New England Biolabs). The plasmid for expression of SPC-TCR variant as transformed into BL21(DE3) and cells were grown at 37 °C to an $OD_{600}$ of 0.6−0.8. Isopropyl β-D-1-thiogalactopyranoside (IPTG, 1 mM) was added, and the cells were collected 3.5-4 h later.

For purification, typically 3-4 g of the cell paste was resuspended with 10 mL of buffer per g of cell paste with Buffer A [50 mM $NaPO_4$ (pH 8.0), 100 mM NaCl, 5% glycerol, and 5 mM BME] supplemented with 5 mM Imidazole, 1 mM PMSF (GoldBio), a protease inhibitor tablet (Thermo Scientific), and 4 kU DNase nuclease (Fisher Scientific). Cells were treated with r-lysozyme (45kU/g of cell paste; MilliPore Sigma) and disrupted via sonication. Extract was clarified by centrifugation and was added to 1.0-1.5 mL of Ni-NTA resin. The column was washed with 50 column volumes (CV) of buffer A with 5 mM imidazole and 10 CV of buffer A with 20 mM imidazole. The protein was eluted with buffer A with 300 mM imidazole. Protein containing fractions were combined and concentrated using Amicon ultracentrifugal filter units (10 kD, MilliporeSigma) and buffer exchanged by a PD10 column (Fisher Scientific) into buffer A without glycerol. The protein was concentrated to 10-30 mg/mL and stored at -80 °C. The concentrations of SPC-fused constructs were determined using a Bradford assay and confirmed by $A_{280}$ using a computationally predicted extinction coefficient (ExPASy). Additionally, SDS-PAGE was used to ensure each variant used in the FA competition assay was of similar purity.

The SNAP-Nar1 construct was generated by replacing the SUMO tag in a 6xHis-SUMO Nar1-10mer plasmid (pTB146) described before (11) with a SNAP-tag using Gibson Assembly. The SNAP-tag insert was obtained from the pSNAP-tag(T7)-2 plasmid (New England Biolabs) by digestion with *NdeI* and *PacI*. Assembly was carried out using a Gibson Assembly Kit (New England Biolabs) and the success of the assembled construct was confirmed by DNA sequencing. Experssion and purification was carried out as described for SPC constructs.

**Pri1 cloning and purification.** The plasmid with full-length *Sc*Pri1 coding sequence in pDONR201 was obtained from the plasmid repository at Arizona State University (DNASU). The Pri1 coding region was inserted into the pDEST17 vector via Gateway cloning to create Pri1 with N-terminal 6xHis-tag ($^{His}$Pri1). The F409A, D408A, and L407A mutants were created via Q5 mutagenesis. For expression, the plasmid was transformed into Rosetta2(DE3) (Novagen) and induced as described for SPC constructs, except the temperature was shifted to 18˚C just before IPTG addition, and cells were harvested 18 h later. Protein was purified as described for SPC except buffer B [50 mM Bis-Tris (pH 6.5), 300 mM NaCl, 10% glycerol, and 5 mM BME] was used in place of buffer A and the column was washed with 50 CV of buffer B with 5 mM Imidazole and 10 CV of buffer B with 20 mM Imidazole. After elution and buffer exchange into buffer B, protein was concentrated to 7-12 mg/mL and stored at -80 ˚C.

**Fluorescence Anisotropy (FA) Equilibrium Binding assay.** The FITC-labeled 4-mer peptide (HLDF or HLDW; GenScript) with N-terminal FITC linked to the peptide via a 1,6-aminohexanoic acid spacer was dissolved in water containing 0.125% ammonium hydroxide. The concentration of the FITC-labeled peptides was determined from its absorbance at 493 nm and the FITC extinction coefficient (70,000 $M^{-1}$ $cm^{-1}$). The assay was carried out and data analyzed as previously described.(12) Briefly, *Ct*Cia1-Cia2 (0-50 µM) was added to either 0.1 µM for FITC-HQDW and FITC-HLDF or 0.05 µM for FITC-HLDW. The observed anisotropy values were plotted against protein concentration and fit to determine $K_D$ value, constraining the $A_{max}$ value to the value observed with FITC-HQDW (180 mA). For presentation, data was normalized as described.(12)

**Competitive Fluorescence Anisotropy (FA) Assays.** Competitive FA assays were performed in 96-well polypropylene black plates (Corning Costar, #3356) at 25˚C. *Ct*Cia1-Cia2 (25 µM for assays using FITC-HQDW or 3-5 µM for assays using FITC-HLDW) was mixed with FITC-HQDW (0.1 µM) or FITC-HLDW (0.05 µM) and 0.1 mg/mL BSA in the Anisotropy Buffer [50 mM $NaPO_4$ (pH 8.0), 100 mM NaCl, 5% glycerol, and 5 mM BME]. After 1 h, unlabeled competitor (SPC-variants, peptides, or small molecules; 0–2 mM) was added so that the final volume in each well was 200 µL. After 1h, the fluorescence anisotropy was measured with a SpectraMax5 plate reader (Molecular Devices) with excitation and emission wavelengths at 488 and 520 nm, respectively, using a 515 nm cutoff filter in high-sensitivity mode (for FITC-HQDW) or low-sensitivity mode (for FITC-HLDW) with 100 reads/well. The anisotropy was calculated and plotted versus the concentration of the unlabeled competitor and f fitted to a hyperbolic competitive inhibitor model (**Equation 1**) in GraphPad Prism 7:

$$A = A_{max} - \left( (A_{max} - A_{min}) \cdot \left( \frac{x}{x + IC_{50}} \right) \right)$$

where $A_{max}$ is the maximum anisotropy, $A_{min}$ is the minimum anisotropy, $x$ is the concentration of the unlabeled competitor, and $IC_{50}$ is the concentration of the unlabeled competitor required for 50% displacement of the FITC-labeled tetrapeptide.

For the TCR motif variant with "LDW" consensus motif (**Fig. S4B** and **S4D**), the data was fitted to the quadratic competitive binding equation (**Equation 2**):

$$A = A_{max} - \left( (A_{max} - A_{min}) \cdot 0.5 \cdot \left( \frac{(x + P + IC_{50}) - \sqrt{(x + P + IC_{50})^2 - 4 \cdot x \cdot P}}{P} \right) \right)$$

where $A_{max}$ is the maximum anisotropy, $A_{min}$ is the minimum anisotropy, $x$ is the concentration of the unlabeled competitor, $IC_{50}$ is the concentration of the competitor required for 50% displacement of the FITC-labeled tetrapeptide, $P$ is the concentration of the *Ct*Cia1-Cia2.

To simplify presentation in figures, the data were normalized using **Equation 3**:

$$A_{norm} = \frac{A_{raw} - A_{f.min}}{A_{f.max} - A_{f.min}}$$

where $A_{f.min}$ and $A_{f.max}$ were determined by fitting the raw data to Equation 1 or 2. To evaluate variants unable to achieve full displacement of the FITC-probe due to their low binding affinities, the minimum anisotropy was constrained during normalization to match the value observed for reference (i.e., wild-type) constructs. The normalized anisotropy was plotted against the protein concentration and fitted to Equation 1 or 2, constraining $A_{max}$ to 1 and $A_{min}$ to 0.

The dissociation constant ($K_d$) for the unlabeled competitor was calculated using the Cheng-Prusoff equation **(Equation 4)**:

$$K_D = \frac{IC_{50}}{\left(1 + \frac{[FITC]}{K_D^{FITC}}\right)}$$

where [*FITC*] is the concentration of the FITC-labeled probe (0.05 μM for FITC-HLDW and 0.1 μM for others), and $K_D^{FITC}$ is the dissociation constant for the FITC-labeled probe-Cia1-Cia2 interaction.

The synthesized peptides, Nar1-10-mer: KDLVSVGSTW and modified Leu1-9-mer: TFDKVHLDW (GenScript), were dissolved in water with 0.125% ammonium hydroxide and quantified using the extinction coefficient (5500 $M^{-1}cm^{-1}$ based on the prediction from ExPASy).

Commercially available amino acids (N-acetyl-W, -F, -Y, and their analogs) were dissolved in the buffer containing 50 mM $NaPO_4$ (pH 8.0) and 100 mM NaCl. The concentration of the amino acid solutions was determined using their reported extinction coefficients: N-acetyl-W and related compounds, 5600 $M^{-1}cm^{-1}$ at 280 nm; N-acetyl-Y, 1400 $M^{-1}cm^{-1}$ at 274 nm; and N-acetyl-F, 200 $M^{-1}cm^{-1}$ at 257 nm.(31)

**Affinity copurification assay.** Protein-protein interaction testing was carried out as described.(11) Briefly, 350 $\mu$g of Cia1-Cia2 was mixed with 1.5-fold excess (~500 $\mu$g) of *Sc*Pri1 and incubated in Assay Buffer [50 mM HEPES (pH 7.0), 150 mM NaCl, 5% Glycerol and 5 mM BME] for 1 h at 4˚C and then chromatographed through streptactin resin (IBA). Input and elution fractions were analyzed by SDS-PAGE with Coomassie staining.

**Bioinformatic analysis.** CIA client protein sequences (n=699) were obtained and analyzed as described.(11, 12) Briefly, the sequences containing C-terminal Trp (88 sequences) and Phe (87 sequences) were analyzed separately. To identify consensus amino acids, the enrichment factor for each amino acid at specific positions was calculated by comparing the observed frequency at those positions with its usage frequency in the human proteome. For visualization, a pie chart or plot was generated to represent the percentage of residues at specific positions with an enrichment factor.

**E-FTMap Analysis.** The E-FTMap algorithm applies a fast Fourier transform (FFT) based computational solvent mapping approach to dock 119 organic probe molecules, sorted into 28 chemically distinct probe sets, within a localized region on a receptor surface.(19) For each probe set, mapping calculations are performed by translating 2000 quasi-uniform rotations of each probe on a 0.8 Å translational grid. From each mapping run, a physics-based scoring function is used to identify energy minima in which each set of probes are predicted to bind. Then, low-energy probe poses from each probe set are selected, superimposed, and clustered on the basis of their chemical features in order to identify pharmacophore regions where hydrogen bond acceptor, hydrogen bond donor, apolar, aromatic, or halogen atoms are predicted to favorably bind. In this study, E-FTMap was applied to the *D. melanogaster (Dm)* Cia1-Cia2 structure (PDB: 6TBN).(10) Mapping calculations were centered on the coordinates of a hot spot identified via the FTMap web server that appeared within a close proximity to experimentally validated binding site residues.(12)

**Generating docked poses of the LDW tripeptide.** Initial conformers of the LDW tripeptide were generated using OpenEye 20.04 Software Suite. In all computations, the N-terminus of the tripeptide was acetylated, and a charged carboxylate was attached to the C-terminus. The classic mode of OMEGA 4.1.2.0 was used to generate 50,000 conformers of the tripeptide.(36) Then the *Dm*Cia1-Cia2 (PDB: 6TBN)(10) was prepared for docking using the "MakeReceptor" graphical utility included in OEDOCKING (version 4.3.1.0, OpenEye Scientific Software, http://www.eyesopen.com).(33-35) A docking box was manually created to encompass residues E105, K107, K125, and R123 on *Dm*Cia1 and Q128 and T123 on *Dm*Cia2b. Next, all conformers of the tripeptide were docked to the prepared receptor using the default settings of the FRED 4.1.1.0 software.(33, 34) The top 600 scoring poses from the docking run were retained for further analysis. Then, the pairwise root mean square distances (RMSD) were calculated between the retained poses, and a simple-greedy clustering algorithm was used to identify cluster centers that had the most neighbors within a 0.5 Å radius. Clusters with fewer than 5 members were removed from consideration, resulting in 21 remaining clusters. These clusters were ranked on the basis of their size. The centroid of the cluster corresponding to Group D was advanced for further study as it was most consistent with the three criteria derived from the biochemical experiments.

**Molecular dynamics simulations.** Molecular Dynamics (MD) simulations were performed using GROMACS 2018.3.(37, 38) The topology of a protein–peptide complex was prepared using parameters from the AMBER99SB force field and TIP3P water model.(40, 41) Systems were solvated using a cubic-box of pre-equilibrated water. The minimum distance between the solute and the edge of the box was defined to be 1.2 nm. To neutralize the system, NA and CL counterions were added to the simulation box. The solvent system was minimized for 10,000 steps using the steepest descent algorithm and then for an additional 100 steps using the conjugate gradient algorithm. During solvent minimization, all solute heavy atoms were restrained using a force constant of 41,840 kJ/mol $nm^2$. Following solvent minimization, protein and peptide side chains were relaxed. The entire system was minimized for 10,000 steps using the steepest descent algorithm, followed by 100 steps of minimization using the conjugate gradient algorithm. During these steps, protein and peptide backbone atoms were restrained using force constant of 2092 kJ/mol $nm^2$.

After minimization, NVT simulations were run to gradually heat the system from 5–300 K over the course of 50 ps. Then, the system underwent 1.0 ns of NPT equilibration, where the pressure was held constant at 1.01325 bar. Following equilibration, production runs were performed in the NPT ensemble for 10–100 ns at a temperature of 300 K and a pressure of 1.01325 bar with a 2 fs time step. Simulation data was recorded every 500 time steps. All simulations were run using the leap-frog stochastic dynamics (SD) integrator as implemented in GROMACS 2018.3.(39) The LINCS algorithm was used to constrain covalent bonds involving hydrogen atoms.(42, 43) Periodic boundary conditions were applied in all three dimensions surrounding the cubic unit cell. Nonbonded interacting pair lists were calculated using the Verlet cutoff scheme, which was updated every 20 time steps. The nonbonded interaction cutoff was set to 1.2 nm. For Van der Waals interactions, the force-switch modifier was used to gradually shift the interaction potential to zero from 1.0–1.2 nm. Long-range electrostatic interactions were calculated using the Particle Mesh Ewald summation with a Fourier spacing of 0.16 nm and cubic interpolation. The SD integrator was used as a thermostat by setting the friction constant to 0.5 $ps^{-1}$. To control pressure, a Berendsen barostat with isotropic pressure coupling, a coupling time constant of 2.0 ps, and an isothermal compressibility of 4.5 x $10^{-5}$ bar was used.(42) During all simulations, protein backbone atoms were restrained using a 2092 kJ/mol $nm^2$ force constant. For each peptide sequence that was studied with MD, 10 independent simulations were initiated using the minimization, equilibration, and production protocols described above.

**AlphaFold analysis.** AlphaFold3 modeling was performed using the available webserver.(17) Cia1, Cia2, and CIA client sequences were obtained from the UNIPROT database. For prediction of *Dm*Cia-Cia2 bound to a TCR peptide, a peptide corresponding to the final 9 residues of *Sc*Leu1 was used, with leucine at the –2 position in place of the glutamine. Figures were generated via Pymol3, with electrostatic surface potential using the APBS electrostatics plugin.(44)
